# Genome editing reveals reproductive and developmental dependencies on specific types of vitellogenin in zebrafish (*Danio rerio*)

**DOI:** 10.1101/456053

**Authors:** Ozlem Yilmaz, Amelie Patinote, Thaovi Nguyen, Emmanuelle Com, Charles Pineau, Julien Bobe

## Abstract

Oviparous vertebrates produce multiple forms of vitellogenin (Vtg), the major source of yolk nutrients, but little is known about their individual contributions to reproduction and development. This study employed a CRISPR/Cas9 genome editing to assess essentiality and functionality of zebrafish (*Danio rerio*) type-I and -III Vtgs. The multiple CRISPR approach employed to knock out (KO) all genes encoding type-I *vtgs* (*vtg1, 4, 5, 6,* and *7*) simultaneously (*vtg1*-KO), and the type-III *vtg* (*vtg3*) individually (*vtg3*-KO). Results of PCR genotyping and sequencing, qPCR, LC-MS/MS and Western blotting showed that only *vtg6* and *vtg7* escaped Cas9 editing. In fish whose remaining type-I *vtgs* were incapacitated (*vtg*1-KO), and in *vtg3*-KO fish, significant increases in Vtg7 transcript and protein levels occurred in liver and eggs, a heretofore-unknown mechanism of genetic compensation to regulate Vtg homeostasis. Fecundity was more than doubled in *vtg1*-KO females, and fertility was ~halved in *vtg3*-KO females. Substantial mortality was evident in *vtg3*-KO eggs/embryos after only 8 h of incubation and in *vtg1*-KO embryos after 5 d. Hatching rate and timing were markedly impaired in *vtg* mutant embryos and pericardial and yolk sac/abdominal edema and spinal lordosis were evident in the larvae, with feeding and motor activities also being absent in *vtg1*-KO larvae. By late larval stages, *vtg* mutations were either completely lethal (*vtg1*-KO) or nearly so (*vtg3*-KO). These novel findings offer the first experimental evidence that different types of vertebrate Vtg are essential and have disparate requisite functions at different times during both reproduction and development.

## 1. INTRODUCTION

In oviparous animals, maternally supplied vitellogenins (Vtgs) are the major source of yolk nutrients supporting early development. Vertebrate Vtgs are specialized members of a superfamily of large lipid transfer proteins that are preferentially produced by the liver and transported via the bloodstream to the ovary (Babin et al., 2007). The Vtgs are taken up into growing oocytes via receptor-mediated endocytosis (Opresko and Wiley, 1987), where they are processed by the lysosomal endopeptidase, cathepsin D, into product yolk proteins that are stored in the ooplasm (Carnevali et al., 1999a, b, 2006). Jawed vertebrates produce three major forms of Vtg arising from a *vtg* gene cluster that was present in the ancestor of tetrapods and ray-finned fish (Babin, 2008; Finn et al., 2009). During vertebrate evolution these ancestral *vtg* genes were subject to whole genome duplications, loss of paralogs and lineage-specific tandem duplications, giving rise to substantial variation in the repertoire and number of *vtg* genes present in an individual species, especially among teleost fish (Andersen et al., 2017). The linear yolk protein domain structure of complete teleost Vtgs is: NH_2_-lipovitellin heavy chain (LvH)-phosvitin (Pv)-lipovitellin light chain (LvL)-beta component (β’c)-C-terminal component (Ct)-COOH (Patiño and Sullivan, 2002; Hiramatsu et al., 2005). Most teleosts possess from two to several forms of A-type Vtg (VtgA), which may be complete or incomplete, as well an incomplete C-type Vtg (VtgC) lacking both Pv and the two small C-terminal yolk protein domains (β’c and Ct). For example, the complex zebrafish (*Danio rerio*) Vtg repertoire includes five type-I Vtgs (Vtg 1, 4, 5, 6 and 7) that are incomplete, lacking β′-c and Ct domains (=ostariophysan VtgAo1), two type-II Vtgs (Vtg2 and Vtg8) that are complete (=VtgAo2), and one type-III Vtg (Vtg3), which is a typical VtgC (Yilmaz et al., 2018).

The multiplicity of teleost Vtgs and the roles that different types of Vtg play in oocyte growth and maturation and in embryonic and larval development has been target of attention for decades (Hiramatsu et al., 2005; Reading and Sullivan, 2011; Sullivan and Yilmaz, 2018). The most diverse group of fishes, the spiny-rayed teleosts (Acanthomorpha) generally possess two paralogous complete forms of VtgA (VtgAa, and VtgAb) in addition to VtgC, and these are orthologs of the zebrafish type-I, type-II and type-III Vtgs, respectively (Finn et al., 2009). In some marine species spawning pelagic eggs, the VtgAa has become neofunctionalized so that its product yolk proteins are highly susceptible to proteolytic degradation by cathepsins during oocyte maturation, yielding a pool of free amino acids (FAA) that osmotically assist oocyte hydration and acquisition of proper egg buoyancy (Matsubara et al., 1999; Finn and Kristoffersen, 2007) and that also serve as critical nutrients during early embryogenesis (Thorsen and Fyhn, 1996; Finn and Fyhn, 2010). The major yolk protein derived from the corresponding VtgAb (LvHAb) is less susceptible to maturational proteolysis. Based on its limited degradation during oocyte growth and maturation, and its utilization late in larval life in some species, it has been proposed that the VtgC may be specialized to deliver large lipoprotein nutrients to late stage larvae without affecting the osmotically active FAA pool (Reading et al., 2009; Reading and Sullivan, 2011). Aside from these few examples, very little is known about specific contributions of the different types of Vtg to developmental processes in acanthomorphs, virtually nothing is known about specialized functions of individual types of Vtg in other vertebrates, and no individual form of Vtg has been proven to be required for the developmental competence of eggs or offspring.

The zebrafish has become an established biomedical model for research on reproduction and developmental biology because they are small, easily bred in the laboratory with short generation time, and lay clutches of numerous large eggs every few days, with external fertilization of the transparent eggs in which embryonic development is easily observed (Ribas and Piferrer, 2013). A reference genome sequence is available, providing the needed databases and bioinformatics tools to conduct genomic and proteomic research on Vtgs in this species. Details on the genomic and protein domain structure of each individual zebrafish Vtg and on their transcript expression and protein abundance profiles were recently made available by Yilmaz et al. (2018). Coupled with these advantages, the presence of multiple genes encoding the three classical major types of Vtg in zebrafish offers a unique opportunity to investigate their essentiality and functionality via application of CRISPR/Cas9 (clustered regularly interspaced short palindromic repeats (CRISPR)/CRISPR-associated protein 9) technology (Doudna and Charpentier, 2014), a powerful gene-editing tool that provides a reliable process for making precise, targeted changes to the genome of living cells.

The extensive multiplicity of genes encoding type-I Vtgs, the major contributors to yolk proteins in zebrafish eggs, is a matter of interest considering their lack of β’c- and Ct- domains, which contain 14 highly conserved cysteine residues that are known to be involved in disulfide linkages required for complex folding of the Vtg polypeptide and possibly for the dimerization of native Vtg thought to be required for binding to its oocyte receptor (Reading et al., 2009; Reading and Sullivan 2011). Additionally, the type-III Vtg (VtgC), lacking all but Lv domains and usually being the least abundant form of Vtg, but one universally present in teleosts, begs investigation regarding its contributions to early development. Therefore, the main objectives of this study were to discover whether type-I Vtgs and type-III Vtg (VtgC) are required for zebrafish reproduction, and to identify specific developmental periods and processes to which they significantly contribute, by investigating the effects of knock out (KO) of their respective genes using the CRISPR/Cas9 gene-editing tool.

## 2. RESULTS

Large deletion mutations of 1821 bp and 1182 bp of gDNA were introduced in zebrafish type-I *vtgs* (*vtg1*-KO) and in *vtg3* (*vtg3*-KO), respectively, via CRISPR/Cas9 genome editing. The introduced deletions involved 703 bp and 714 bp of the respective transcripts, encoding 234 aa and 239 aa of their respective polypeptide sequences, and they resulted in double strand breaks in the ORF in both cases (**Fig 1**, **S1 Fig**). For both *vtg1*-KO and *vtg3*-KO, the introduced mutation altered the structure of the deduced LvH chain of the Vtg polypeptide and, in the case of *vtg3*-KO, it extended into the Vtg receptor-binding domain (**Fig 2**, **S1 Fig**). Introduced mutations were detected by genotyping via PCR screening of gDNA at each generation using combinations of primers flanking each altered target site (**Fig 1**). F0 generation individuals exhibiting a heterozygous mutant double banding pattern were retained as founders for production of stable mutant lines (**Fig 3**).

**Fig 1.**
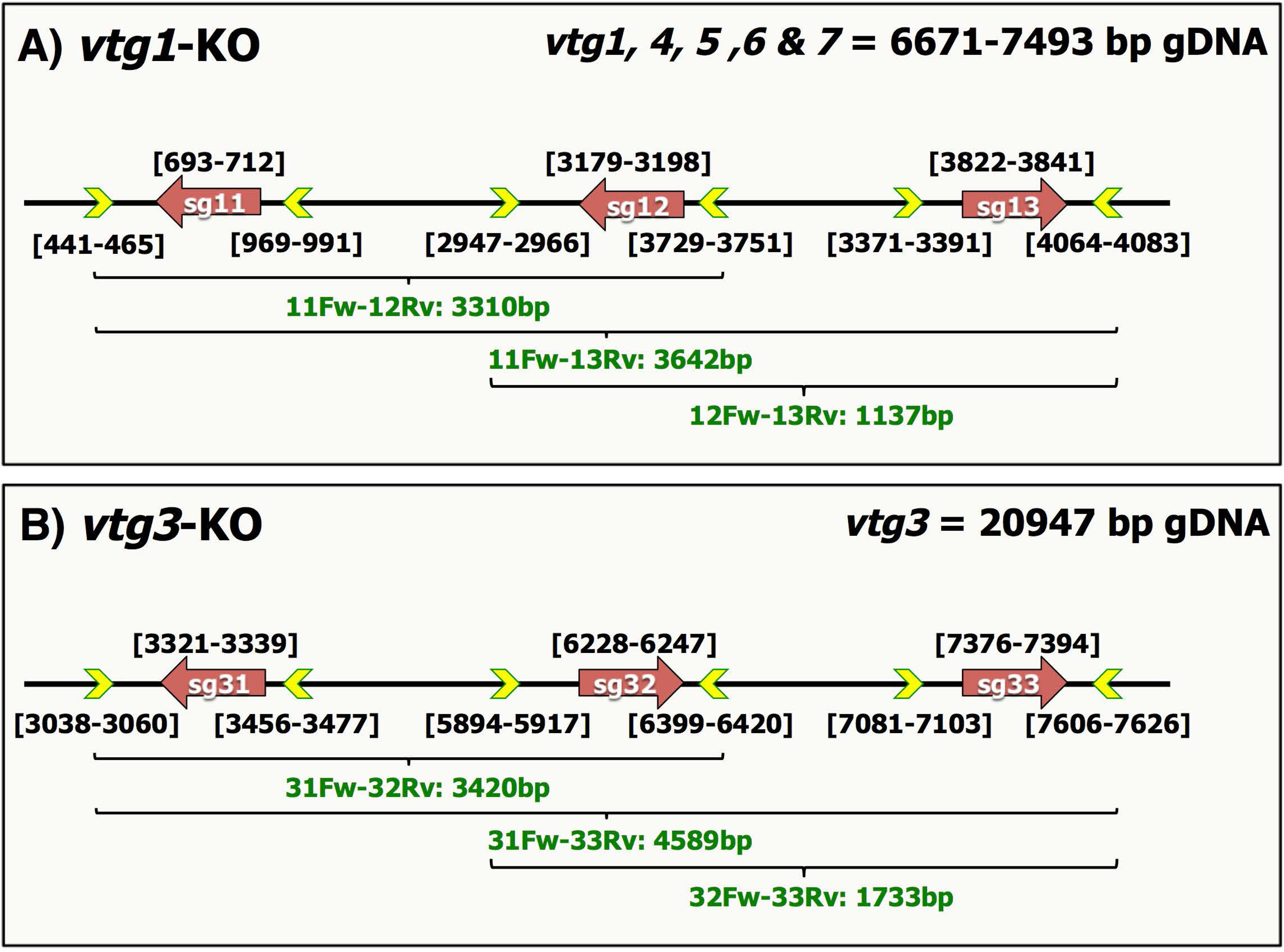
Schematic representation of the general strategy for CRISPR target design in the zebrafish *vtg* knock out (KO) study. **A)** Type-I *vtg* knock out (*vtg1*-KO). *vtg1* is depicted as representative of the five targeted type-I zebrafish *vtg* genes. **B)** Type-III *vtg* knock out (*vtg3*-KO). Target sites are shown by brown colored arrows labeled as sg followed by 1, 2 or 3 indicating the targeted zebrafish *vtg* type and the number of the target site (i.e. sg11, sg12, and sg13: single guide RNAs (sgRNAs) for target sites 1, 2, and 3 for *vtg1,* respectively. sg31, sg32, and sg33: sgRNAs for target sites 1, 2, and 3 for *vtg3,* respectively). Arrows are oriented to indicate the sense/antisense orientation of each target. Numbers above each target site specify its exact location by nucleotide in the genomic sequence of the zebrafish *vtgs.* Primers used in screening for introduced mutations by PCR are shown as yellow arrowheads outlined in green, which are oriented to indicate the sense/antisense orientation of the primer. Numbers below each primer site indicate its exact position by nucleotide in the genomic sequence of the targeted gene (see also **S1 Fig**). Horizontal brackets below indicate areas screened for mutations by PCR using selected primer combinations; bold green text below the brackets indicates the primer pair followed by the size of the band (bp) expected for wild type gDNA in agarose gel electrophoresis (see **Fig 3**). 11Fw, *vtg1* target1 forward primer; 12Rv, *vtg1* target2 reverse primer; 13Rv, *vtg1* target3 reverse primer; 12Fw, *vtg1* target2 forward primer; 31Fw, *vtg3* target1 forward primer; 32Rv, *vtg3* target2 reverse primer; 33Rv, *vtg3* target3 reverse primer; 32Fw, *vtg3* target2 forward primer. Primer sequences are given in **S1 Table**.

**Fig 2.**
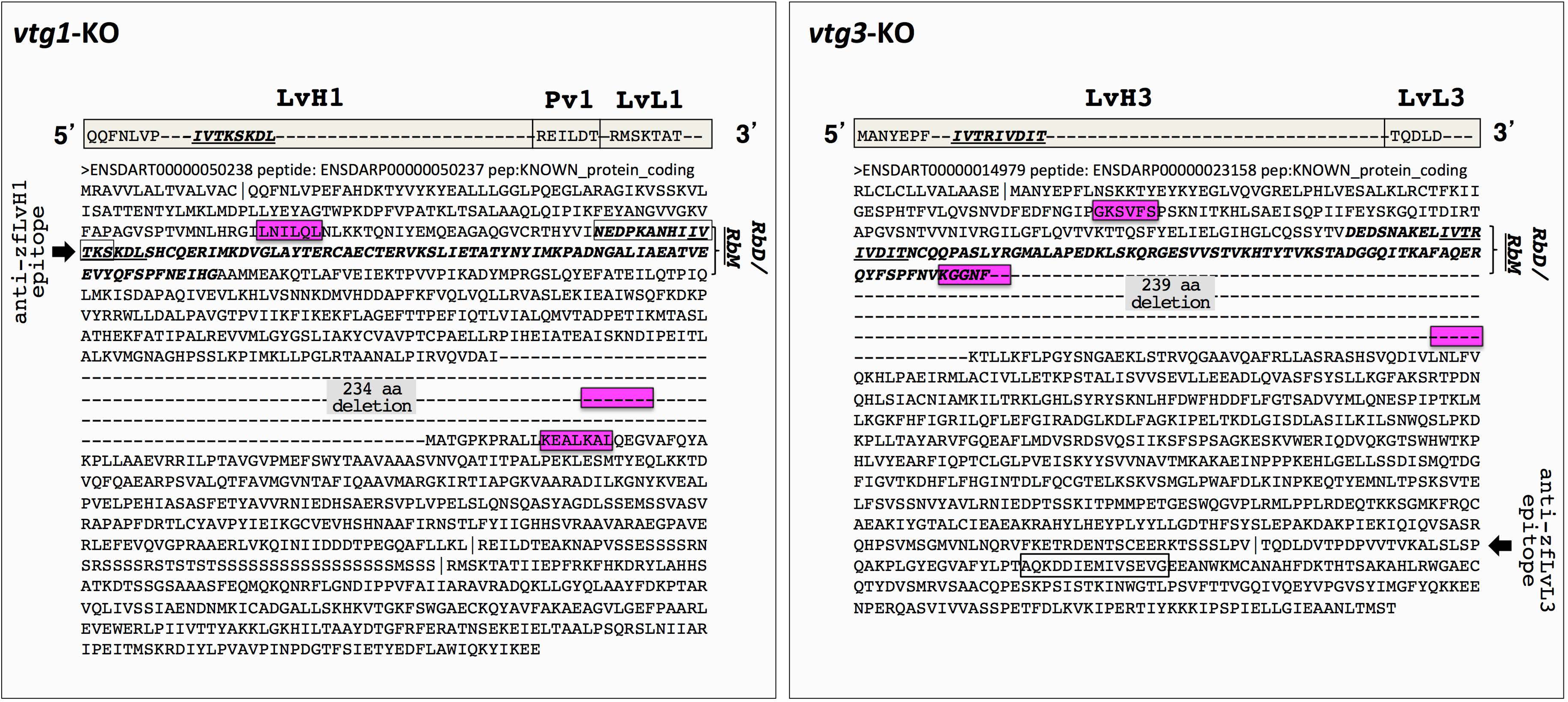
Location of mutations introduced by CRISPR/Cas9 in the predicted polypeptide sequence of targeted zebrafish *vtgs.* The yolk protein domain structures of Vtg1 (representative of zebrafish type-I Vtgs) and Vtg3 are pictured in 5’ > 3’ orientation above each panel. Light gray horizontal bars represent the lipovitellin heavy and light chain (LvH, LvL) and phosvitin (Pv) domains of the respective Vtg (Vtg3 lacks a Pv domain) and are labeled above in large bold type. Sequences within these bars indicate the N-terminus of each yolk protein domain, the starting points of which are also indicated by vertical bars in the polypeptide sequence shown below. The 85-residue Vtg receptor-binding domain ***(RbD)*** and the critical 8-residue Vtg receptor-binding motif ***<u>(RbM)</u>*** located within this domain, which were identified by Li et al. (2003) in the LvH domain of blue tilapia *(Orechromis aureus)* VtgAb, are shown in the polypeptide sequences in boldface italic type, with the ***RbM*** sequence being additionally underlined and also shown in the yolk protein domain map above. Residues encoded by nucleotide sequences targeted by sgRNAs for Cas9 editing are framed in magenta-shaded boxes. Cas9 created mutations (large deletions) are indicated with dashes replacing amino acid (aa) residues and the size of deletions in aa (234 aa and 239 aa for *vtg1*-KO and *vtg3*-KO, respectively) in these regions are labeled by gray shaded text. Short sequences that were employed as epitopes to develop Vtg domain-specific antibodies against Vtg1-LvH (anti-LvH1) and Vtg3-LvL (anti-LvL3) are indicated by framed text on the LvH and LvL domains of Vtg1 and Vtg3, respectively, with their location also highlighted by black arrows labeled with the epitope names given by vertically-oriented text in the panel margins.

Microinjection efficiency was acceptable and high, resulting in 20% and 80% mutation positive embryos at 24h screening, for *vtg1*-KO and *vtg3*-KO, respectively. This efficiency was confirmed by finclip genotyping when these embryos reached adulthood. However, mutation transmission to F1 offspring was as low as 0.010% for *vtg1*-KO and 0.025% for *vtg3*-KO, and only 2 heterozygous (Ht: *vtg1-/+* and *vtg3*-/+) adult males were available to continue reproductive crosses with non-related wild type (Wt: *vtg1*+/+ and *vtg3*+/+) females for production of F2 generations. The rate of mutation transmission to the F2 generation produced from F1 Ht males and Wt females was 55% and 70% for *vtg1*-KO and *vtg3*-KO, respectively. Reproductive crosses of Ht males and Ht females revealed a Mendelian inheritance pattern with 25% wild type (wt: sibling wild type; *vtg1*+/+ and *vtg3*+/+), 52% heterozygous and 22% homozygous (Hm: *vtg1*-/- and *vtg3*-/-) individuals at the F3 generation. Hm F3 females and males were crossed to produce the F4 generation yielding 100% homozygous offspring carrying only the mutated allele (**Fig 3**). As these Hm individuals are generally inviable *(see below),* production of subsequent generations of mutants requires crossbreeding of heterozygotes.

**Fig 3.**
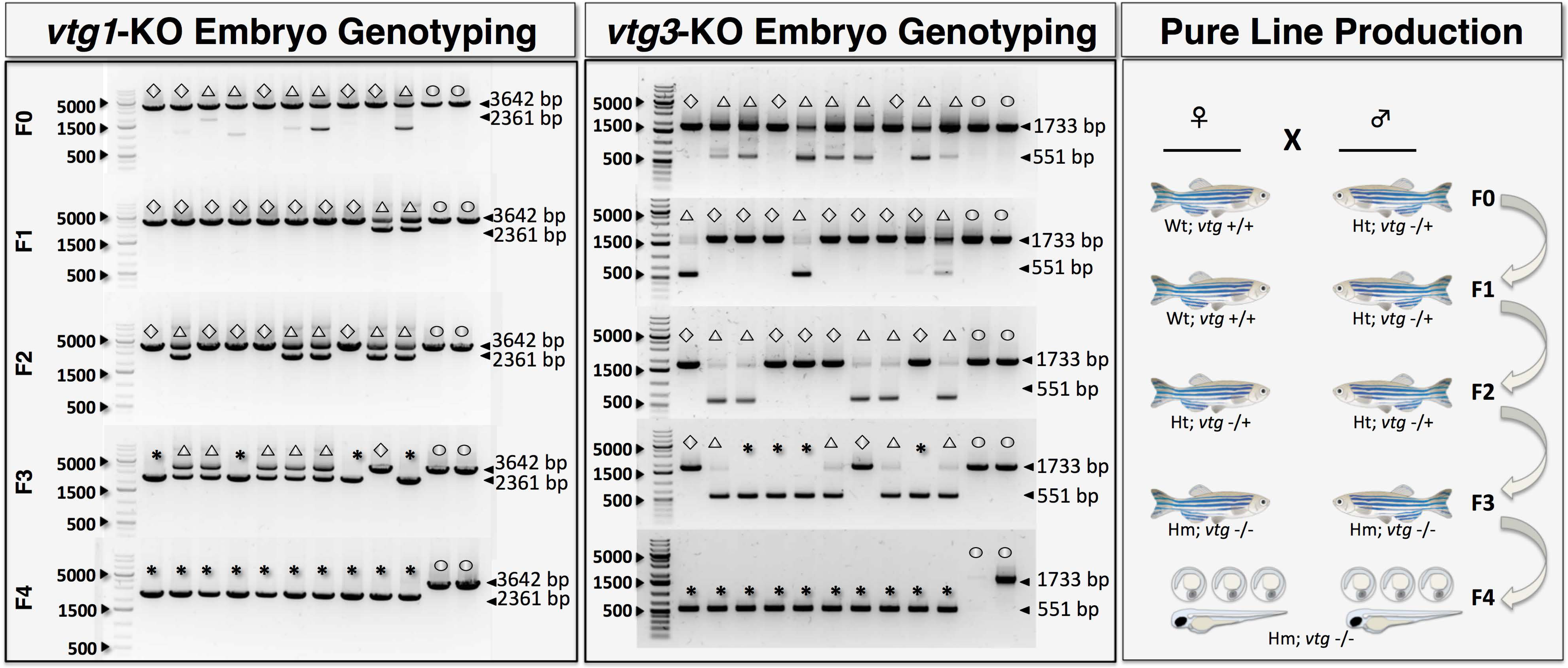
Detection of CRISPR/Cas9-introduced mutations by embryo genotyping and production of F4 generation *vtg*-KO mutants. Left and middle panels illustrate genotyping of embryos at 24 h post-fertilization (hpf) by PCR for *vtg1-KO* (representative of zebrafish type-I *vtgs)* and *vtg3-KO* lines, respectively, from the F0 to F4 generation. F0 indicates the generation reared from microinjected embryos and F1-4 represent offspring raised from each subsequent generation. The agarose gel electrophoresis results shown here represent screening of 10 randomly sampled embryos as representatives of their generations and two additional wild type embryos as controls. Bands comprised of wild type intact gDNA (3642 bp and 1733 bp for *vtg1* and *vtg3,* respectively) and mutated gDNA (2361 bp and 551 bp for *vtg1-*KO and *vtg3-KO,* respectively) are shown and highlighted by black arrowheads on the right side of each panel. Open circles; non-related wild type fish (Wt) carrying intact *(vtg1*+/+ or *vtg3*+/+) genomic DNA. Open diamonds; sibling wild type individuals, which do not carry the desired mutation in either allele *(vtg1*+/+ or *vtg3*+/+) of their gDNA. Open triangles; heterozygous (Ht) individuals carrying the introduced mutation on only a single allele *(vtg1*-/+ or *vtg3*-/+) in their genomic DNA. Asterisks; homozygous embryos (Hm) carrying the introduced mutation in both alleles *(vtg1*-/- or *vtg3*-/-) of their genomic DNA. The panel on the far right illustrates the general strategy followed to establish pure zebrafish lines bearing the desired Cas9 introduced mutation. This process involved stepwise reproductive crosses (indicated by X) between males (♂) and females (♀) indicated here with zebrafish icons. F0-4 represents the zebrafish generations produced in the process. Images of sub-adult fish are shown for simplicity at generation F4; all or most of these fish were actually inviable and did not survive past early developmental stages (see text for details).

For both *vtg* KO lines, the relative level of expression of each individual *vtg* transcript in livers of Hm, Ht, and wt F3 generation females were compared to those obtained for Wt female liver. KO of type-I *vtgs* resulted in the absence of *vtg1, vtg4,* and *vtg5* transcripts in F3 Hm *vtg1*-KO female liver, representing a significant decrease in levels of these transcripts compared to Ht, wt and Wt females (p<0.05). Levels of *vtg6* and *vtg7* transcripts were still detectable, with *vtg7* transcript levels being significantly higher (~3-fold) in Hm *vtg1*-KO female liver as compared to Wt female liver (p<0.05). The *vtg1*-KO had no significant effect on *vtg2* and *vtg3* expression (**Fig 4**, **Panel A**). No *vtg3* transcripts were detected in F3 Hm *vtg*3-KO female livers, representing a significant decrease in *vtg3* transcript levels compared to Ht, wt and Wt females (p<0.05). The F3 Hm *vtg*3-KO females showed a statistically significant ~3-fold increase in hepatic *vtg7* transcript levels relative to Wt fish (p<0.05). No significant effect of *vtg3*-KO on expression of other *vtg* genes was observed (**Fig 4**, **Panel B**).

**Fig 4.**
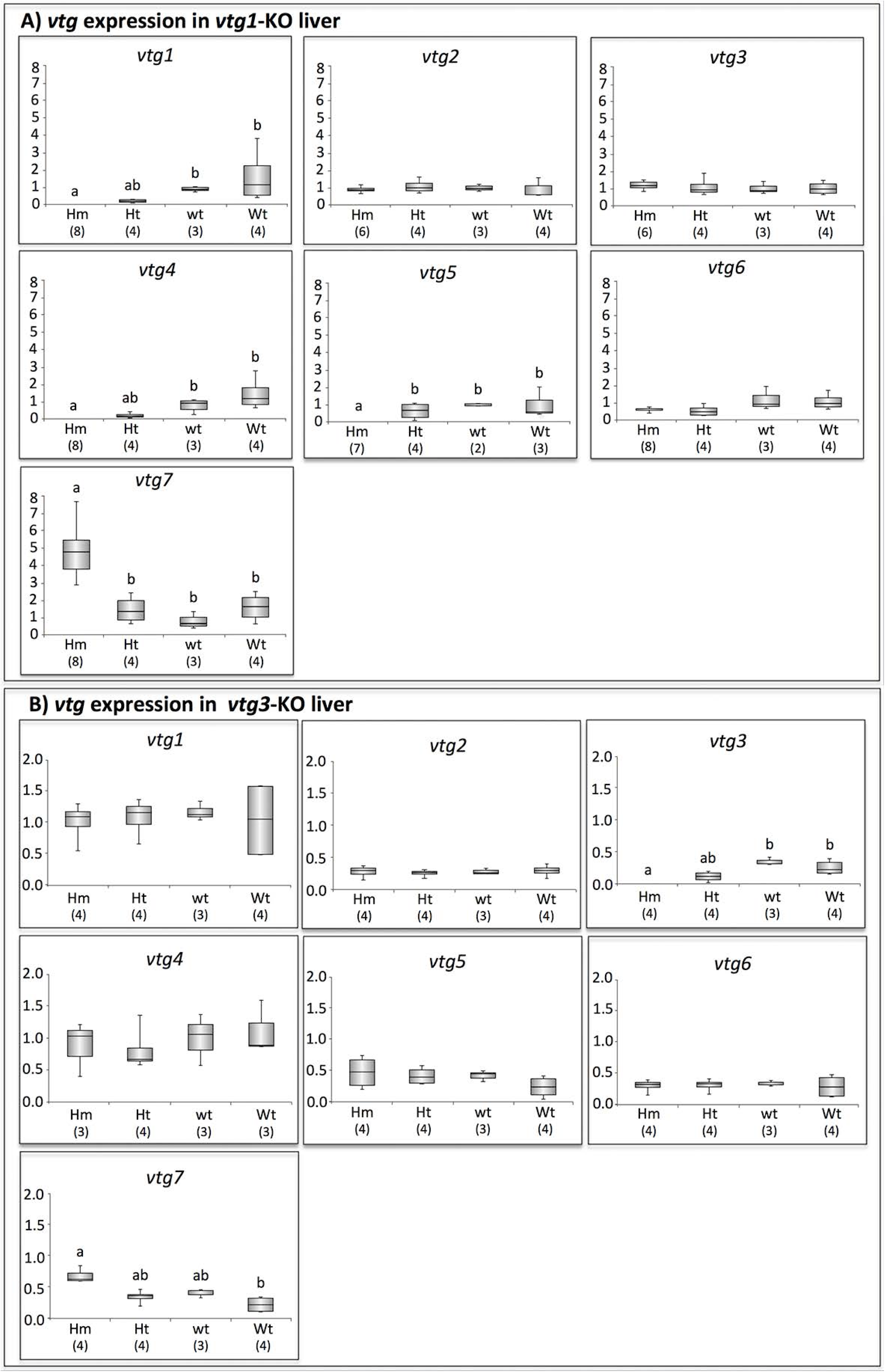
Relative quantification of *vtg* gene expression in *vtg*-KO zebrafish female liver. **A)** Comparison of gene expression levels for all *vtgs* in F3 *vtg1-KO* female liver (Hm, homozygous; Ht, heterozygous; wt, sibling wild type) versus non-related wild type female liver (Wt). TaqMan qPCR-2^-ΔΔCT^ mean relative quantification of gene expression was employed using zebrafish 18S ribosomal RNA *(18S)* as the reference gene. Data were statistically analyzed using a Kruskal Wallis nonparametric test p<0.05 followed by Bejamini Hochberg corrections for multiple tests p<0.1. **B)** Comparison of gene expression levels for all *vtgs* in F3 *vtg3-KO* female liver (Hm, Ht and wt) versus Wt female liver. SYBR Green qPCR-2^-ΔΔCT^ mean relative quantification of gene expression normalized to the geometric mean expression of zebrafish elongation factor 1a *(eif1a),* ribosomal protein L13a *(rpl13a)* and *18S* was employed. Data were statistically analyzed using a Kruskal Wallis nonparametric test p<0.05 followed by Benjamini Hochberg corrections for multiple tests p<0.1. In the box plots, the centerlines indicate the median for each data set, upper boxes indicate the difference of the 3^rd^ quartile from the median, lower boxes indicate the difference of the 1^st^ quartile from the median. Top whiskers indicate difference of the maximum value from the 3^rd^ quartile and the bottom whiskers indicate the difference of the minimum values from the 1^st^ quartile in each data set. In both panels, numbers below x-axis labels indicate sample size and lowercase letters above the error bars represent significant differences between means (p<0.05). For box plots sharing a common letter superscript, the means are not significantly different.

The relative abundances of individual Vtgs or of their product yolk proteins in liver and eggs, respectively, of F3 Hm *vtg1*-KO females were evaluated as normalized spectral counts (N-SC) from LC-MS/MS and revealed no detectable amount of Vtg1, 4 and 5 (p<0.05) (**Fig 5A**). Similar to gene expression levels in these same samples, Vtg6 and 7 protein levels were still detectable and the Vtg7 levels were significantly higher in Hm *vtg1*-KO female liver and eggs than in corresponding samples from Wt females. (p<0.05). The relative abundance of Vtg7 protein was ~4-fold and ~3-fold higher in Hm *vtg1*-KO liver and eggs, respectively, than in Wt females. Additionally, even though they were uniformly low, Vtg3 protein levels were also significantly higher (~2-fold) in Hm *vtg1*-KO eggs than in Wt eggs (p<0.05) (**Fig 5A**).

**Fig 5.**
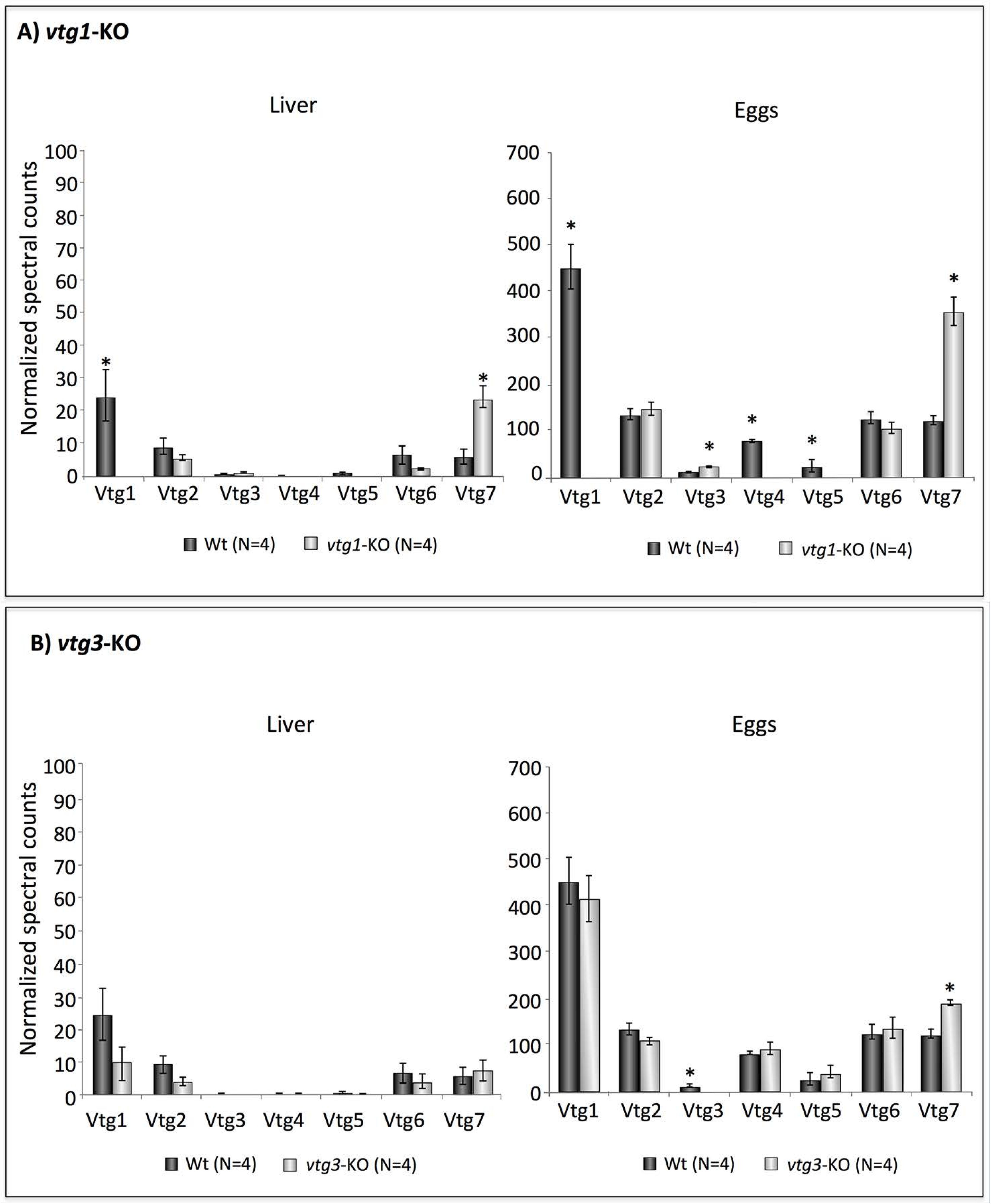
Relative quantification of multiple vitellogenins by LC-MS/MS in *vtg*-KO zebrafish female liver and eggs. **A)** Comparisons of mean normalized spectral counts (N-SC) for Vtg protein levels in Wt versus Hm F3 *vtg1*-KO female zebrafish livers and in eggs obtained from these females, indicated by dark and light gray vertical bars, respectively. Vertical brackets indicate SEM. **B)** Corresponding comparison of N-SC for Vtg protein levels in Wt versus F3 Hm *vtg3-KO* female zebrafish livers and in eggs obtained from these females. Asterisks indicate statistically significant differences between group means detected by an independent samples Kruskal Wallis non-parametric test (p<0.05) followed by Benjamini Hochberg correction for multiple tests (p<0.1)

The *vtg3*-KO resulted in the absence of detectable Vtg3 protein in both liver and eggs of F3 Hm *vtg3*-KO females (p<0.05) but it did not seem to influence the relative abundances of Vtg1, 2, 4, 5, and 6 or their yolk protein products in these samples (**Fig 5B**). However, Vtg7 protein levels were significantly higher (~1.5-fold) in *vtg3*-KO eggs than in Wt eggs (p<0.05), but *vtg3*-KO did not significantly alter the relative abundance of Vtg7 protein in the liver of the egg donors, although average levels were higher for the *vtg3*-KO fish (**Fig 5** **Panel B**). For *vtg1*-KO, *vtg3*-KO and Wt females, relative protein abundances of all detected Vtgs were generally lower in liver in comparison to eggs. Among the various forms of Vtg protein, their relative abundance in eggs from Wt females ranged from 15 to 31 times higher than in livers of the same fish.

Domain-specific, affinity purified polyclonal antibodies were developed in rabbits against zebrafish (zf) Vtg Type-specific epitopes (Type-I Vtg: NEDPKANHIIVTKS on LvH1; Type-III Vtg: AQKDDIEMIVSEVG on LvL3. See **Fig 2**). The antibodies were used to detect these proteins by Western blotting in the respective Hm *vtg*-KO, Ht and Wt female livers, ovaries and eggs. The rabbit anti-zfLvH1 antibody revealed the presence of high molecular weight bands corresponding in mass to LvH1 in all tested individuals and tissues (*data not shown*), consistent with the reported escape of the *vtg6* and *vtg7* from Cas9 editing and the presence of Vtg6 and Vtg7 protein in liver, ovary and eggs from all groups of fish in the *vtg1*-KO experiment (Hm, Ht and Wt). In Western blots performed using anti-zfLvL3 in the *vtg3*-KO experiment, the antibody detected mainly a bold ~24 kDa band in samples of both ovary and eggs from Ht and Wt fish, but not from Hm fish, very close to the deduced mass of the LvL3 polypeptide (21.3 kDa) (Yilmaz et al., 2018) (**Fig 6**). The distinct absence of this ~24 kDa band only in samples of Hm ovary and eggs is considered to be evidence of successful *vtg3*-KO in this experiment. The very bold ~68 kDa band also present in samples of ovary and eggs from Ht and Wt females, but absent in samples from Hm *vtg3*-KO fish, which have a faint band in this position, may represent a degradation product of intact, covalently linked LvH-LvL conjugate (Vtg3) persisting after maturational proteolysis, as has been described for several species (Reading et al., 2009). Faint high molecular weight bands mainly ≥ 68 kDa were also evident for samples of liver, ovary and eggs from all groups of fish in the *vtg3*-KO experiment (Hm, Ht and Wt). For Hm *vtg3*-KO fish these bands are taken to indicate slight cross-reactivity of the antibody with yolk proteins other than LvL3 under the experimental conditions employed. For the corresponding Ht and Wt fish, some of these bands may represent high molecular weight Vtg3 products bearing intact or partially degraded LvL3, as noted above. No bands specific to Ht and Wt fish were detected in Western blots of liver performed using this antibody, consistent with absence of significant quantities of Vtg3 protein detectable in Wt liver by LC-MS/MS (**Fig 5**), a commonly observed phenomenon (see Yilmaz et al. (2016)) suggesting that Vtg3 is rapidly released into the bloodstream after synthesis.

**Fig 6.**
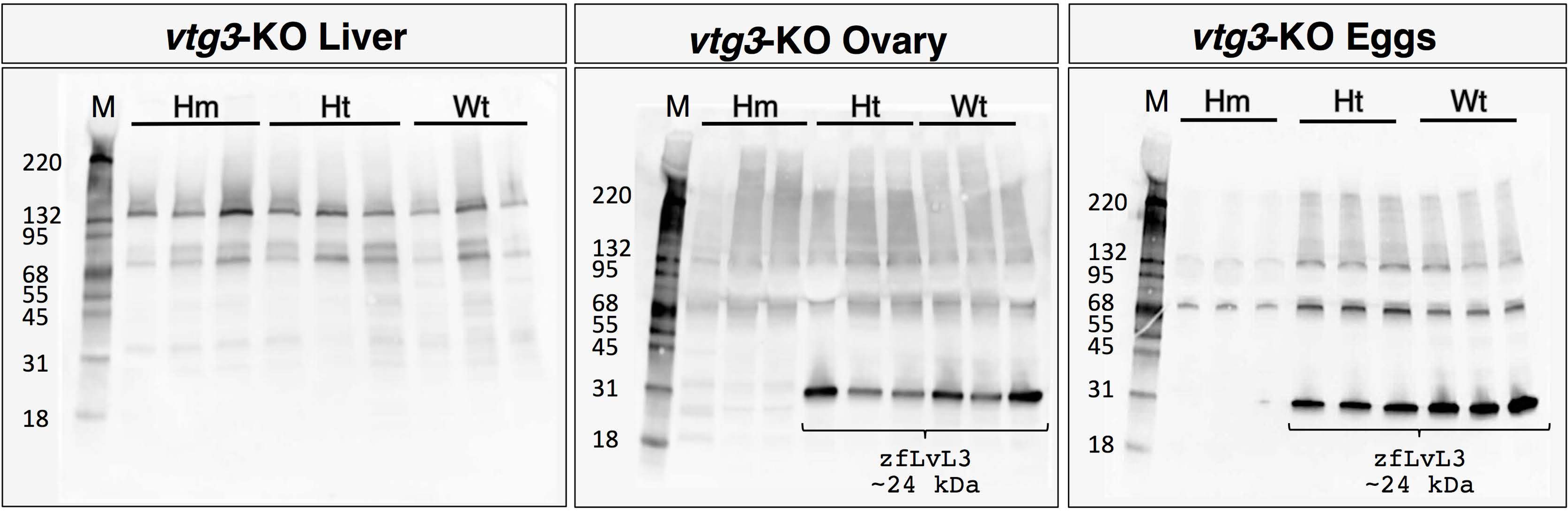
Detection of Vtg3 in *vtg3*-KO versus wild type female liver, ovary and eggs by Western blotting. An affinity-purified, polyclonal anti-zfLvL3 antibody was employed to detect LvL3 in this experiment. Numbers on the left of each panel indicate the mass of molecular weight marker proteins (kDa). M, marker protein ladder; Hm, homozygous; Ht, heterozygous; Wt, non-related wild type. Bands that were detected in Ht and Wt zebrafish whose mass corresponds to that of zebrafish Vtg3 LvL (LvL3, ~24 kDa) are indicated with brackets and labels immediately underneath (LvL3).

Phenotypic parameters including fecundity (number of eggs per spawn), egg fertilization, hatching and survival rates, and egg diameter (embryo and chorion diameter) as well as larval size at 8 days post spawning (dps), were measured to detect potential effects of *vtg* KO on zebrafish reproductive performance and development. There were no significant differences between Hm *vtg1*-KO and Wt eggs or offspring in fertilization rate, embryo size or larval size, respectively (**Fig 7**). However, F3 Hm *vtg1*-KO females produced significantly more eggs per spawn (593 ± 40.06, mean ± SEM) than did Wt females (280 ± 28.97) (p<0.05), although the final hatching rate of these eggs at 10 dps (64.9 ± 6.45 %) was significantly lower than for eggs from Wt females (99.6 ± 0.24 %) (p<0.05). Eggs from F3 Hm *vtg1-*KO females also were strikingly delayed in hatching, completing hatching at 9 dps versus 5 dps for control fish (**Fig 7**). It was noted that the Hm *vtg1*-KO embryos appeared to have weaker heartbeats and body movements during incubation antecedent to hatching as compared to *vtg3*-KO embryos, which, even with malformations, exhibited apparently normal heartbeat rhythms and body movements comparable to those seen in Wt embryos. Embryo and larval survival rates of Hm *vtg1*-KO offspring were also significantly lower than for Wt offspring, beginning from 5 dps when their mean survival rate was 57.14 ± 7.34 % compared to 79.40 ± 5.75 % for Wt fish. The survival rate of Wt offspring changed little thereafter, whereas the survival rate of Hm *vtg1*-KO offspring continued to decline, with *vtg1*-KO being completely lethal to the larvae by 16 dps (**Fig 8**).

**Fig 7.**
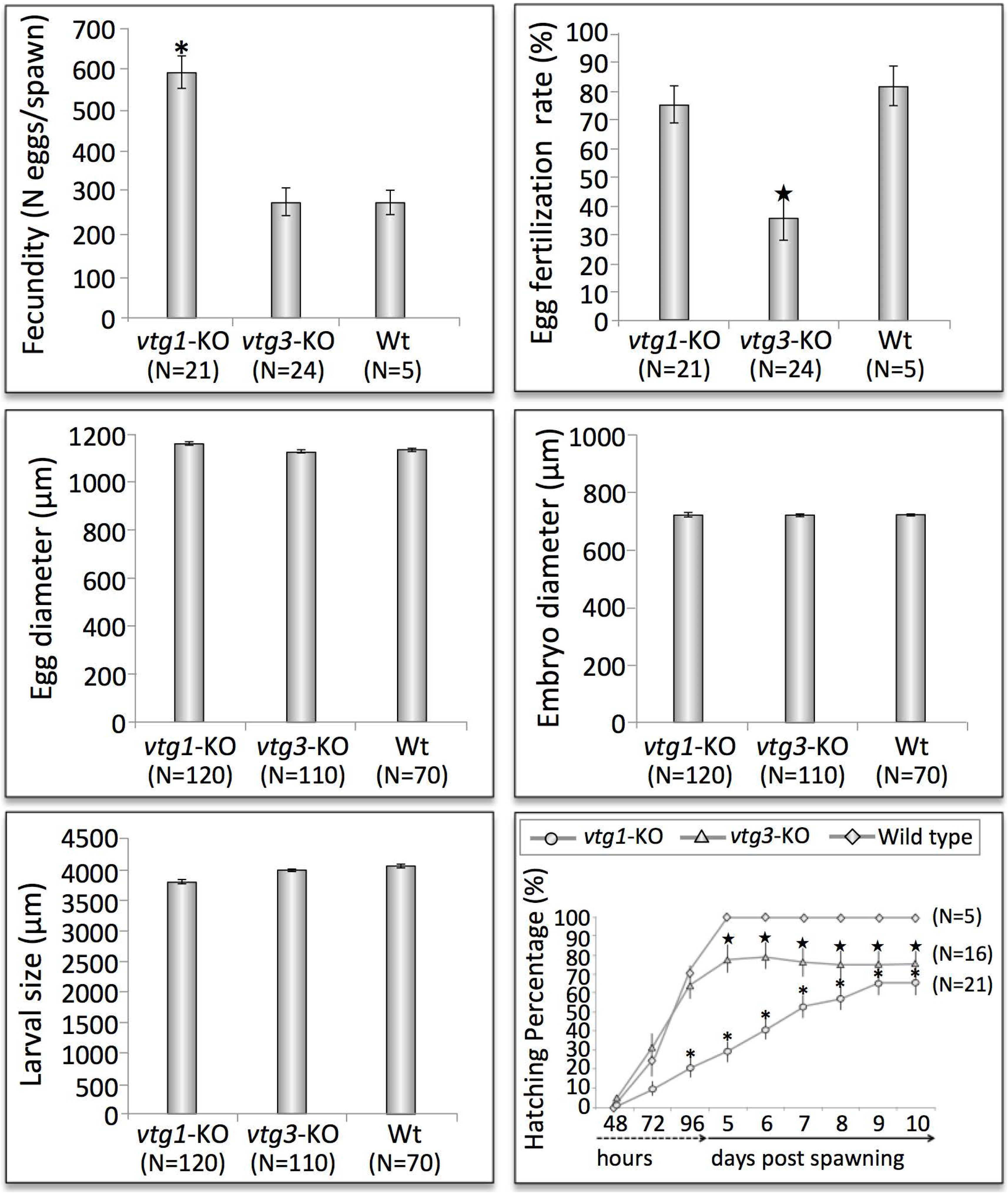
Phenotypic measurements of F3 *vtg*-KO females and their F4 progeny. Bar graphs indicate mean values (±SEM) for measurements of each parameter and labels below the x-axes indicate the groups that were compared. In the panel at the bottom right, mean hatching percentages for Hm *vtg1*-KO, Hm *vtg3*-KO, and Wt eggs are shown as circles, triangles and diamonds, respectively. Numbers on the x-axis accompanied by dashed- and solid-lined arrows represent sampling times in hours or days post spawning, respectively. In all graphs, asterisks and black stars indicate mean values that are statistically significantly different from corresponding Wt mean values based upon results of an independent samples t-test (p<0.01) followed by Benjamini Hochberg corrections for multiple tests in the case of hatching percentage (p<0.05).

**Fig 8.**
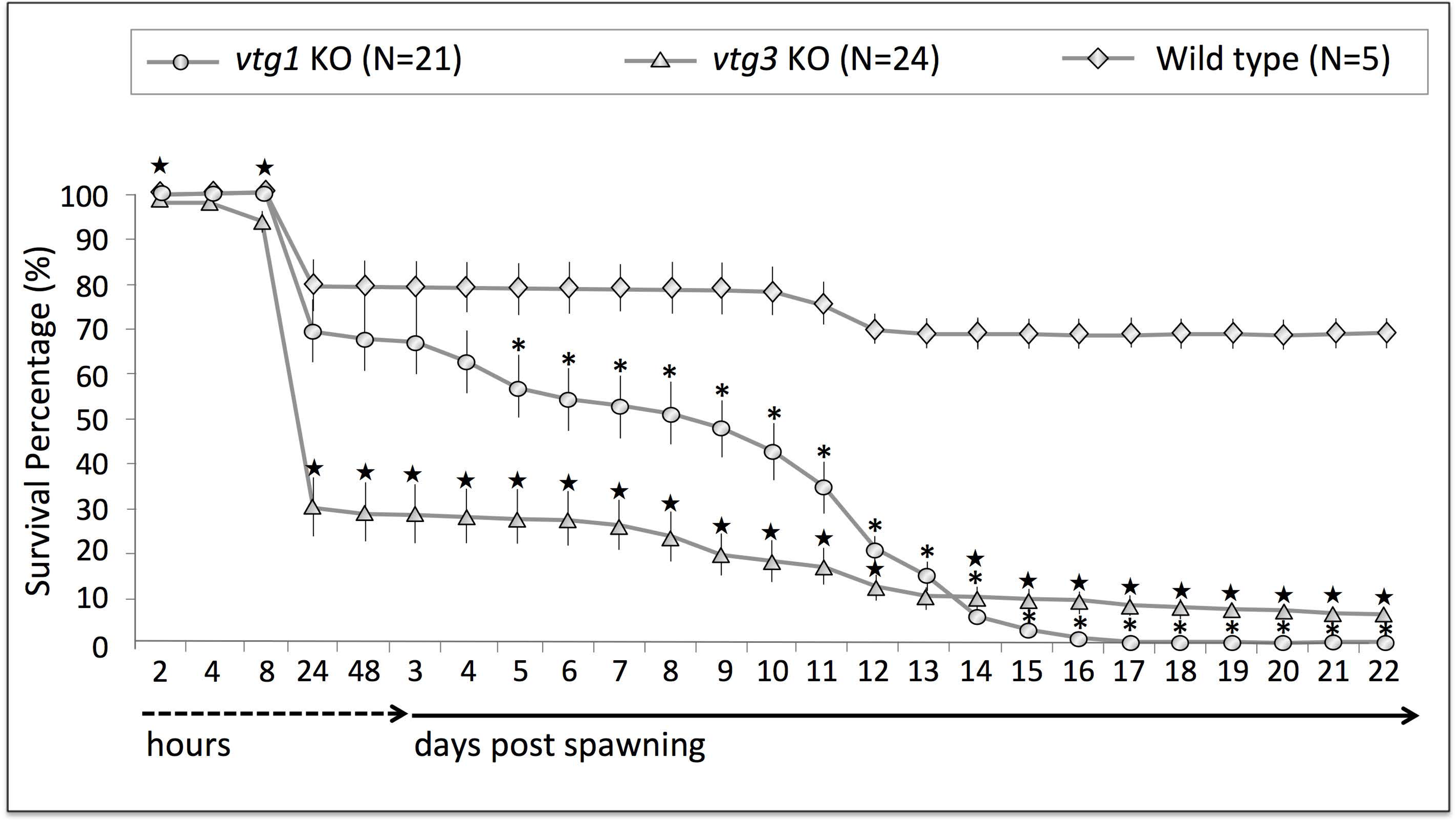
Comparisons of survival percentages for homozygous F4 *vtg1*-KO and *vtg3*-KO zebrafish embryos and larvae versus wild type offspring. Line plots represent mean survival percentages and numbers on the x-axis accompanied by dashed- and solid-lined arrows represent sampling times in hours or days post spawning during the observation period. Mean survival percentages for Hm *vtg1*-KO, Hm *vtg3*-KO and unrelated Wt embryos and larvae at each time point are indicated by circles, triangles and diamonds, respectively, and vertical lines indicate SEM. Asterisks and black stars indicate mean values that are statistically significantly different from corresponding mean wild type (Wt) values based upon results of an independent samples t-test (p<0.01) followed by Benjamini Hochberg corrections for multiple tests (p<0.05).

There were no significant differences between Hm *vtg3*-KO fish and Wt fish in fecundity, embryo size or larval size (**Fig 7**). However, the fertility, hatching rate and overall survival of Hm *vtg3-*KO eggs and offspring, respectively, were significantly less than seen in Wt fish (p<0.05) (**Fig 7**). The fertilization rate of eggs from F3 Hm *vtg3*-KO females (35.5 ± 7.7 %) was substantially lower than for Wt eggs (81.6 ± 7.0 %), although hatching of eggs from these females was only slightly delayed, and to a much lesser extent than was observed for eggs from the F3 Hm *vtg1*-KO females (*see below*). The final hatching rate for eggs obtained from F3 Hm *vtg3*-KO females was 74.3 ± 7.7 % at 10 dps compared to 99.6 ± 0.24 % for Wt eggs (**Fig 7**). Embryo and larval survival rates of Hm *vtg3*-KO offspring were significantly less than for Wt offspring (p<0.05), beginning from 8 hours post spawning (hps), with the difference from Wt fish increasing throughout the 22 d experiment (**Fig 8**). As previously reported (Yilmaz et al., 2017), at 2–4 hps eggs from low fertility spawns have a high incidence of abnormal embryos with asymmetric cell cleavage and/or developmental arrest at early cleavage stages. Such embryos may survive to 8 hps but not to 24 hps. The larval survival rate for Hm *vtg3*-KO offspring was only 6.25 ± 1.6 % at 22 dps compared to 69.2 ± 3.8 % for Wt offspring (**Fig 8**).

Separate panels in **Fig 9** illustrate morphological disorders observed during development of F4 Hm *vtg1*-KO and Hm *vtg3*-KO fish in comparison to offspring from Wt females at 4 and 8 dps. In Hm *vtg*-KO fish, these phenotypic disorders mainly involved pericardial and yolk sac/abdominal edema accompanied by spinal lordosis evidenced as curved or bent back deformities. The severity of these malformations, mainly the pericardial and yolk sac edema, appeared to be relatively lower in Hm *vtg1*-KO fish than in Hm *vtg3*-KO fish. However, the prevalence of deformity was much greater for Hm *vtg1-*KO fish, with nearly all larvae exhibiting some deformity versus approximately 30 % of Hm *vtg1*-KO larvae. Finally, the Hm *vtg1*-KO larvae exhibited no feeding activity or motor activities comparable to those seen in Hm *vtg3*-KO and Wt fish at the same times.

**Fig 9.**
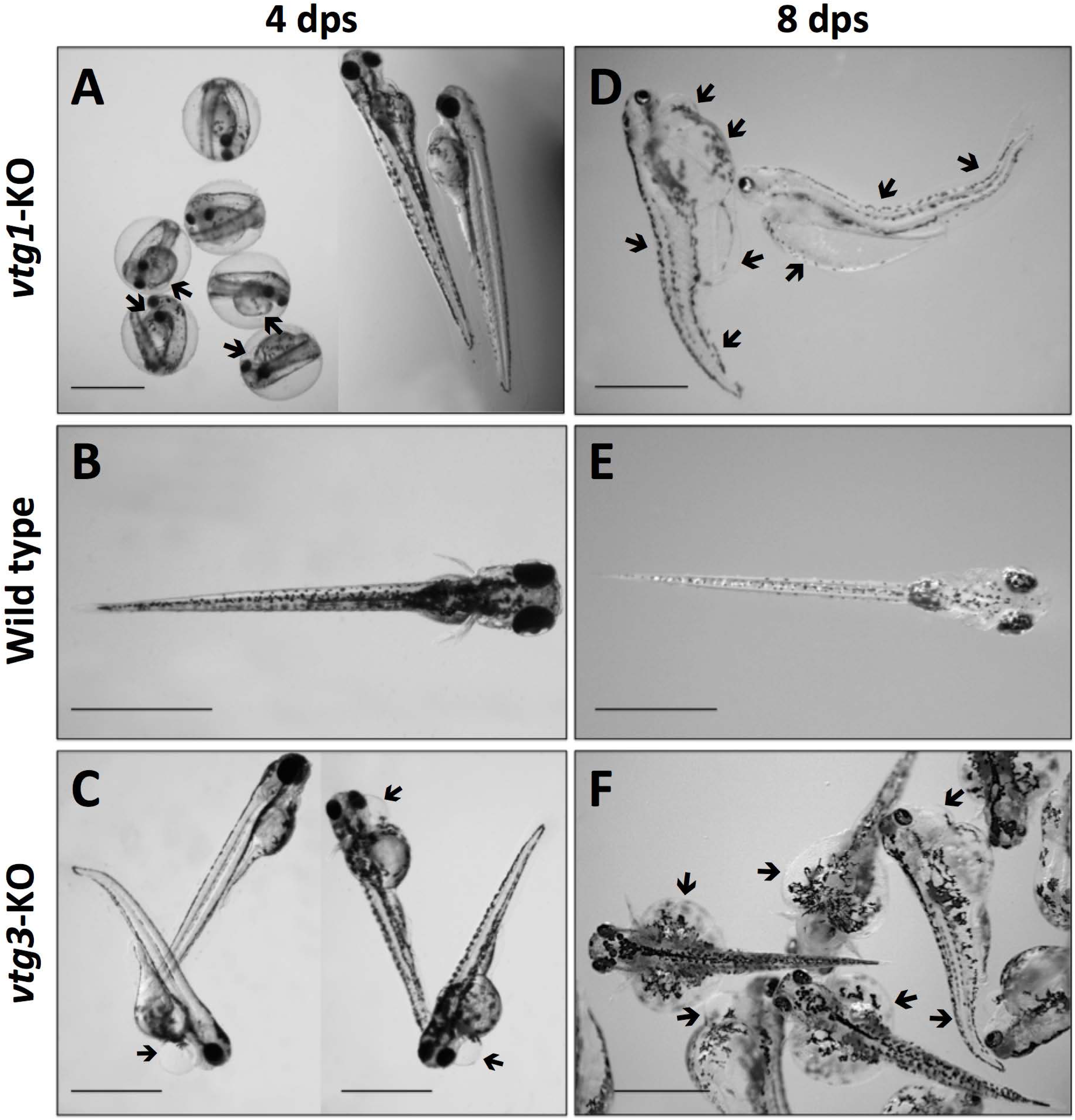
Observed phenotypes of F4 *vtg*-KO offspring compared to wild type offspring. **A) Hm** *vtg1*-KO unhatched embryos and hatched larvae at 4 dps. **B)** Wt larva at 4 dps. **C)** Hm *vtg3*-KO larvae at 4 dps. **D)** Hm *vtg1*-KO larvae at 8 dps. **E) Wt** larva at 8 dps. **F) Hm** *vtg3*-KO larvae at 8 dps. Special features of observed phenotypes are indicated by pointed arrows (see text for details). In all images, horizontal bars indicate 1000 µm.

## 3. DISCUSSION

Vitellogenins are the ‘mother proteins’ that supply most yolk nutrients supporting early vertebrate development, and most species have evolved multiple forms of Vtg. However, little is known about specific functions of these different forms of Vtg and it is uncertain which forms are essential for successful development or at what stage(s) of development they are required. The present research was undertaken to address these questions using a zebrafish CRISPR/Cas9 *vtg* gene KO model. Three out of five type-I zebrafish *vtg* genes (*vtg1, 4* and *5*) were knocked out simultaneously (*vtg1*-KO experiment), and the type-III *vtg* gene (*vtg3*) was knocked out individually (*vtg3*-KO experiment), and the effects on maternal reproductive physiology and offspring development and survival were evaluated.

The efficacy of CRISPR/Cas9, which is reported to be the most practical and efficient tool available for genome editing, was lower in the *vtg1*-KO experiment (20 %), where five genes were targeted concomitantly, than in the *vtg3*-KO experiment (80 %) where only a single gene was targeted. Since the site-specific cleavage efficiency is mostly dependent on the concentrations of single guide (sg) RNAs and Cas9 endonuclease, Liu et al. (2018) related the low efficiency of simultaneous knockout of multiple homologous genes to the fact that more sgRNAs and gene target sites share the same Cas9 enzyme. In addition to low efficiencies in the *vtg1*-KO experiment, the escape of the type-I *vtg6* and *vtg7* from Cas9 editing might be simply an outcome of an insufficient amount of administered Cas9 RNA. Attempts at optimization of sgRNA/Cas9 concentrations may be useful in future studies. Taking into account the syntenic organization and close proximity of type-I *vtg* genes in zebrafish (Yilmaz et al., 2018), and the identity (100 %) of the common target sites for these genes, it is difficult to postulate criteria upon which any preference of Cas9 activity might be directed. No matter which gene-editing tool is used, low efficiency of germline mutant transmission has been a commonly faced problem among researchers, usually leading to labor intensive and time consuming screening work to acquire high-throughputs (Xie et al., 2016). The low ratios of mutation transmission to next generations in the present study (0.01 % - 0.025 %) emphasize the need for further research to improve germline transmission efficiencies in genome editing. Production of stable mutant lines was delayed an extra generation in both the *vtg1*-KO and *vtg3*-KO experiments since no mutation-positive female founders were obtained at the F1 generation for performing subsequent reproductive crosses.

The incapacitation of *vtg1, vtg4,* and *vtg5* in the *vtg1*-KO experiment, and of *vtg3* in the *vtg3*-KO experiment, was confirmed by conventional PCR, agarose gel electrophoresis and sequencing of gDNA and also by relative quantification of corresponding *vtg* transcript and Vtg protein abundances via qPCR and LC-MS/MS, respectively, with the absence of Vtg3 in Hm *vtg3*-KO ovary and eggs being additionally confirmed by Western blotting. Collectively, these procedures provided strong evidence of the success of genome editing and *vtg* gene KO. The introduced mutations were all large deletions (1182-1281 bp) that were achieved by administration of multiple sgRNAs. By disturbing the structure of the LvH chain in both the *vtg1*-KO and *vtg3*-KO experiments via the creation of large gaps in the respective LvH polypeptides, it was expected that the mutant proteins would not fold properly, or be able to bind to their receptor in the case of Vtg3, even if they were produced and partly expressed by the liver. There were no signs of hepatic synthesis of Vtg1, 4 or 5 in Hm *vtg1*-KO individuals or of Vtg3 in Hm *vtg3*-KO liver. While detection of the *vtg6* and *vtg7* transcripts and their product proteins was expected in the *vtg1*-KO experiment, since these two type-I *vtg* genes escaped Cas9 editing, the strikingly high abundance of Vtg7 (but not Vtg6) at both transcript and protein levels in both Hm *vtg1*-KO and Hm *vtg3*-KO individuals (**Figs 4** **and** **5**) was unexpected. These observations suggest an attempt of the organism to compensate for the loss of other types of Vtgs by augmentation of Vtg7 levels, and they imply the existence of heretofore-unknown mechanisms for regulating Vtg homeostasis.

The lack of a mutant phenotype in Hm mutant individuals due to compensatory gene expression triggered upstream of protein function is known as ‘genetic compensation’ and this phenomenon has been encountered in gene editing studies of a wide range of model organisms. As examples, Marschang et al. (2004) related the normal development and lack of mutant phenotypes in LDL receptor-related protein 1b (*LRP1b*)-deficient mutant mice to functional compensation by *LRP1,* and Sztal et al. (2018) found that a genetic *actin1b* (*actc1b*) zebrafish mutant exhibits only mild muscle defects and is unaffected by injection of an *actc1b*-targeting morpholino due to compensatory transcriptional upregulation of an *actin* paralog in the same fish. In the present study, compensatory increases in relative levels of total Vtg protein attributable to upregulation of Vtg7 protein in F4 Hm *vtg1*-KO eggs offset only about half of the decrease in total Vtg protein attributable to KO of *vtg1, 4* and *5* (**Fig 5A**). Therefore, these eggs/offspring were still deficient of type-I Vtg protein and they uniformly exhibited mutant and ultimately lethal phenotypes, perhaps due to the insufficient compensation. In contrast, the compensatory increase in total Vtg protein attributable to upregulation of Vtg7 in Hm *vtg3*-KO eggs was several-fold greater than the loss of Vtg protein attributable to *vtg3* KO (**Fig 5B**), yet many of these eggs/offspring still exhibited mutant phenotypes, with egg fertility being very low (*see below*) and most offspring not surviving for 22 d of development. Nonetheless, the incidence of mutant phenotypes in Hm *vtg3*-KO larvae (30 %) was far less than in Hm *vtg1*-KO larvae, all of which were malformed, and a low percentage (6.25 %) of Hm *vtg3*-KO larvae did survive for 22 d post fertilization, whereas no Hm *vtg1*-KO larvae did. These observations indicate that, while it is possible that upregulation of Vtg7 may have mitigated to some extent the effects *vtg3* KO owing to decreased total Vtg protein, Vtg7 cannot fully substitute for Vtg3 or eliminate the adverse effects of *vtg3* KO on egg fertility and offspring development. Therefore, Vtg3 must have functional properties distinct from Vtg7 and perhaps other type-I Vtgs.

Transcription of *vtg* genes is initiated when estrogen (E2)/estrogen receptor (Esr) complexes bind to estrogen response elements (ERE) located in the gene promoter regions (Babin, 2008; Nelson and Habibi, 2013). E2-Esr complexes can also be tethered to transcription factor complexes targeting binding sites distinct from EREs, and several transcription factors other than Esrs have binding sites located in promoter regions of zebrafish *vtg* genes (reviewed by Lubzens et al., 2017). There is evidence that the multiple *vtg* genes in zebrafish exhibit differential sensitivities to estrogen induction as well as disparate patterns of ERE and other transcription factor binding sites in their promoter regions (Levi et al., 2009, 2012). Bioinformatics analyses indicated that the promoter region of *vtg7* is comparatively rich in binding sites for transcription factors involved in retinoic acid signaling such as retinoic acid response elements (RAREs), and peroxisome proliferator-activated receptors (PPARs)/ retinoid X receptor (RXR), while having only a single ERE (most other *vtgs* having 2-3) (see Levi et al., 2012 Table 3). These types of differences between *vtg* promoters could underpin selective upregulation of Vtg7 in response to ablation of other forms of Vtg (other type-I Vtgs, Vtg3) via gene KO. Conspecific Vtg (type not specified) has been shown to downregulate plasma levels of E2 *in vivo* when injected into vitellogenic rainbow trout (*Oncorhynchus mykiss*) (Reis-Henriques et al., 1997) and to inhibit steroidogenesis leading to E2 production *in vitro* by ovarian follicles of rainbow trout (Reis-Henriques et al., 1997, 2000) and greenback flounder, *Rhombosolea tapirina* (Sun and Pankhurst, 2006). Partial release from such inhibition in *vtg-*KO fish would increase vitellogenic signaling to the liver, activating estrogen responsive genes including those encoding Vtgs, Esr (Esrs) and PPARs.

Whether Vtg7 itself has vitellogenic properties remains to be determined. Certain conspecific Vtgs have been shown to upregulate vitellogenesis in Indian walking catfish (*Clarias batrachus*) (Juin et al., 2017; Bhattacharya et al., 2018) and comparisons of the available deduced catfish Vtg polypeptide sequences (85 and 152 residues; Juin et al., 2017, Fig. 3) to Vtgs from zebrafish and other teleosts using CLUSTAL W and BLASTP (*data not shown*) indicate that they are forms of VtgAo1 showing a high identity to type-I zebrafish Vtgs (up to 80% with Vtg7). The specific mechanism(s) by which Vtg7 is preferentially upregulated in *vtg*-KO zebrafish, and special properties of Vtg7 for regulation of Vtg homeostasis, are meaningful subjects for future research. Levels of Vtg3 protein were also upregulated in eggs from F3 Hm *vtg1*-KO females (**Fig 5A**) but the significance of this increase is difficult to interpret as it was too slight to have much impact on total Vtg levels, and because hepatic levels of *vtg3* transcripts and of Vtg3 protein were not elevated in these fish (**Figs 4A** **and** **5A**). Transcripts of *vtg3* are reported to be the most intensely upregulated transcripts in vitellogenic female and estrogenized male zebrafish (Levi et al., 2009) and there may not have been scope for further increases in the *vtg1*-KO fish. In this case post-transcriptional mechanisms for upregulating Vtg3 could have been at play (Flouriot et al., 1996; Ren et al., 1996). As noted above, Vtg3 may be released into the bloodstream immediately after synthesis, which would explain the lack of significant quantities of this protein in livers of Hm *vtg1*-KO and Wt fish (**Fig 5**).

Neither *vtg1*-KO nor *vtg3*-KO influenced egg, embryo or larval size in spawns producing F4 offspring of the stable mutant lines (**Fig 7**), and there were no apparent differences in ovary structure among the different groups of maternal F3 females (Hm, Ht, wt and Wt) sampled after spawning (*data not shown*). However, F3 Hm *vtg1*-KO females exhibited a 2-fold increase in fecundity (egg production) relative to Wt females, with normal egg fertility equivalent to that of Wt females (**Fig 7**). This response to elimination of three type-I Vtgs (including the most abundant one, Vtg1) implies that one or more of these Vtgs are normally involved in restriction of fecundity, perhaps via the aforementioned inhibition of follicular estrogenesis. It is also possible that Vtg7, which was highly elevated in Hm *vtg1*-KO females, might somehow positively modulate fecundity. The referenced VtgAo1 of walking catfish, when pelleted and implanted into pre-vitellogenic females, has been shown to stimulate vitellogenesis and complete oocyte growth all the way through the transition to final maturation (Bhattacharya et al., 2018). In the final analysis, any ‘compensation’ by Vtg7 for loss of other type-I Vtgs must be deemed ineffectual, as the resulting embryos unconditionally exhibited serious and lethal developmental abnormalities (*see below*).

The *vtg*-KO zebrafish larvae exhibited major phenotypic disorders, mainly pericardial and yolk sac/abdominal edemas and spinal lordosis associated with curved or arched back deformities. These abnormalities were observed to be much less prevalent, albeit usually more severe, in *vtg3*-KO larvae, but present to some extent in all *vtg1*-KO larvae along with the noted behavioral differences. Skeletal axis malformations and pericardial and yolk sac/abdominal edema are among the most common deformities observed in cultured teleosts and they form an interrelated cluster of abnormalities that tend to be observed together (Alix et al., 2017). For example, in zebrafish pericardial edema tends to precede development of yolk sac edema, which when severe leads to notochord deformation (see Hanke et al., 2013, Fig. 1). These abnormailites have been associated with a broad variety of conditions including, as examples, rearing systems for Eurasian perch, *Perca fluviatilis* (Alix et al., 2017), larval rearing temperatures for Atlantic halibut, *Hippoglossus hippoglossus L.* (Ottesen and Bolla, 1998), embryo cryopreservation practices for streaked prochilod, *Prochilodus lineatus* (Costa et al., 2017), and, in zebrafish, phenanthroline toxicity (Ellis and Crawford, 2016), influenza A virus infection (Gabor et al., 2014), knockdown or KO of genes related to kidney function or development, respectively (Hanke et al., 2013; Zhang et al., 2018), knockdown of the *wwox* tumor suppressor gene (Tsuruwaka et al., 2015), deletion of a gene (*pr130*) encoding a protein essential for myocardium formation and cardiac contractile function (Yang et al., 2016), and mutagenesis of genes involved in thyroid morphogenesis and function (Trubiroha et al., 2018), among others. The edemas may ultimately result from many different proximal causes such as cardiac, kidney, liver or osmoregulatory failure, and researchers are just beginning to develop screens to differentiate between them (Hanke et al., 2013). Although they can occur under many different conditions and arise via several possible mechanisms, these major mutant phenotypes observed in the present study were not encountered in control Wt offspring and, therefore, they are clearly related to deficiencies of type-I Vtgs (Vtg1, 4 and 5) and of Vtg3.

Embryo and larval survival rates were severely diminished by *vtg* gene KO, but the magnitude, type and timing of losses differed between *vtg1*-KO and *vtg3*-KO fish (**Fig 8**). The fertility of Hm *vtg3-*KO eggs was only half that observed in Wt eggs (**Fig 7**), indicating that Vtg3 is an important contributor to fertility in zebrafish. Among the Vtgs examined, this dependency was specific to Vtg3, since fertility was not ‘rescued’ by the increase in Vtg7 levels in Hm *vtg3*-KO eggs, which was far greater than normal Vtg3 levels in Wt fish (**Fig 5B**), and this adverse effect on fertility was not seen in Hm *vtg1*-KO eggs. The substantial losses of Hm *vtg3*-KO eggs began early, at only 8 hps, and less than 30% survived to 24 hps (**Fig 8**). Both *vtg1*-KO and Wt eggs showed significant but much fewer losses (p<0.05) during this same interval. In this study, fertility was estimated conservatively, based on numbers of viable embryos showing normal cell division and subsequently developing to ~24 hps. It is uncertain whether the high mortality of Hm *vtg3*-KO eggs between 8 and 24 hps (**Fig 8**) resulted from a failure to be fertilized or from defects in early development involving zygotes that fail to initiate cell division or that briefly undergo abnormal cell divisions and then die. In future studies, some Hm mutant and Wt females should be bred with males bearing a unique germline marker gene, such as *vasa*::*egfp* (Krøvel and Olsen, 2002), that can be genotyped in resulting eggs and embryos to resolve this question.

The mechanism(s) whereby Vtg3 deficiency impairs fertility and/or early development of zebrafish are unknown. A recent study examining the proteomics of egg/embryo developmental competence in zebrafish identified disruption of normal oocyte maturation, including maturational proteolysis of Vtgs, as a likely cause of poor egg quality (Yilmaz et al., 2017). The proteolysis of Vtgs by cathepsins during oocyte maturation, a phenomenon that has been observed in zebrafish eggs undergoing maturation *in vitro* (Carnevali et al., 2006), releases FAA that steepen the osmotic gradient driving water influx through aquaporins on the cell surface, leading to oocyte hydration (Cerdà et al., 2007, 2013). These FAA are also major substrates for aerobic energy metabolism during early embryogenesis (Thorsen and Fyhn, 1996; Finn and Fyhn, 2010). In some species, Vtg3 (VtgC) is subjected to maturational proteolysis (see Yilmaz et al., 2016) and it is possible that zebrafish Vtg3 contributes to these critical processes ongoing during oocyte maturation, which are required for production of viable eggs. However, mass balance considerations seem to exclude the possibility that the early mortality of Hm *vtg3*-KO embryos results substantially from nutritional deficiencies. In this and prior studies of zebrafish, Vtg3 has been shown to be a very minor form of Vtg making only a miniscule contribution to stores of Vtg-derived yolk proteins in eggs (**Fig 5**; see also Yilmaz et al., 2018). Nonetheless, Vtg3 is clearly an important, if not essential, contributor to fertility and/or early development in zebrafish. The continuous mortality of Hm *vtg3*-KO embryos after 24 hps, leading to only ~6% survival at 22 dps, suggests that Vtg3 also contributes to late embryonic and larval development, as suggested in several prior studies (see below).

Survival of embryos emanating from Hm *vtg1*-KO females remained relatively high at 24 hps (~70%) but decreased continuously thereafter, becoming significantly less than survival of Wt embryos by 5 dps, and then decreasing to zero by 16 dps (**Fig 8**). The collective absence of Vtg1, 4 and 5 in zebrafish is lethal to offspring, and this effect could not be rescued via genetic compensation by Vtg7 or offset by the remaining intact Vtgs. This finding is not surprising as, collectively, these 3 type-I Vtgs account for the vast majority of Vtg-derived protein in Wt zebrafish eggs (**Fig 5**; see also Yilmaz et al., 2018). Since most mortality of *vtg1*-KO offspring occurred relatively late in development in larvae, with mortality rate increasing after 10 dps when yolk sac absorption was being completed (**Fig 8**), the collective contributions of Vtg1, 4 and 5 to survival could be largely nutritional, although this remains to be verified.

It is evermore apparent that the different types of vertebrate Vtg can have dissimilar effects on reproductive processes. As noted above (see also Introduction), in marine acanthomorphs spawning pelagic eggs in seawater the different types of Vtg can play disparate roles in oocyte hydration, acquisition of egg buoyancy, and early versus late embryonic and larval nutrition (Matsubara and Koya, 1997; Matsubara et al., 1999, 2003; Reith et al., 2001; Sawaguchi et al., 2005, 2006a, 2006b; Finn, 2007). The type-specific ratios of circulating Vtgs (e.g. VtgAa:VtgAb:VtgC) may vary considerably during oocyte growth, but ratios of their derived yolk protein products present in eggs tend to be fixed and characteristic of species (Hiramatsu et al., 2015; Reading et al., 2017). This is also the case in zebrafish as evidenced by the similarity of Vtg profiles by type (and subtype) in Wt fish in the *vtg1*-KO and *vtg3-*KO experiments, and also in comparison to Wt fish in an earlier study (Yilmaz et al., 2018). It is thought that Vtg type-specific ratios of yolk proteins in eggs are maintained via activity of selective receptors for each type of Vtg, which target their specific ligand(s) into different compartments where their yolk protein products undergo disparate degrees of proteolysis during oocyte maturation. The initial abundance and degree of proteolysis of the yolk proteins determines their relative contribution(s) to oocyte hydration, egg buoyancy, FAA nutrition of early embryos and lipoprotein nutrition for late stage larvae (Hiramatsu et al., 2015; Reading et al., 2017). The collective findings of the present study introduce a new point of view on the roles that multiple vitellogenins can play in vertebrate reproduction. Distinctively from what has been reported previously, the present study presents a mixed model of Vtg functionality covering both maternal reproductive physiology and early development of offspring, where type-I Vtgs regulate fecundity and make essential contributions to embryonic morphogenesis, hatching and larval kinesics and survival (Vtg1, 4 and 5), and also provide some homeostatic regulation of total Vtg levels (Vtg7), while Vtg3 (a typical VtgC) is critically important to fertility and early embryogenesis and also influences later development.

In summary, the present study, for the first time, targeted multiple forms of Vtgs for KO at family level using CRISPR/Cas9 technology in the zebrafish, a well-established biomedical model. The collective knock out of *vtg1*, *4*, and *5* and the individual knock out of *vtg3* were achieved successfully. A compensatory increase in *vtg7* at both transcript and protein levels was observed in both types of *vtg* KO mutants. However, this compensation was not effective in rescuing the serious developmental impairments and high mortalities resulting from ablation of three other type-I Vtgs or of Vtg3. By far the most abundant forms of Vtg in zebrafish, the type-I Vtgs appear to have essential developmental and nutritional functions in both embryos and larvae. In spite of being a very minor form of Vtg in zebrafish and most other species, and also the most divergent form, Vtg3 contributes importantly to the developmental potential of zygotes and/or early embryos. Finally, Vtgs appear to have previously unreported regulatory effects on the physiology of maternal females, including limitation of fecundity (type-I Vtgs) and maintenance of fertility (Vtg3). These novel findings represent the first steps toward discovery of the specific functions of multiple vertebrate Vtgs via genome editing. Further physiological studies are necessary to pinpoint the exact molecular mechanisms disturbed in the *vtg* mutants.

## 4. MATERIAL AND METHODS

### 4.1. Animal care, spawning and phenotypic observations

Zebrafish of the Tübingen strain originally emanating from the Nüsslein-Volhard Laboratory (Germany) were obtained from our zebrafish facility (INRA UR1037 LPGP, Rennes, France). The fish were ~15 months of age and of average length ~5.0 cm and average weight ~1.4 g. The zebrafish were housed under standard conditions of photoperiod (14 hours light and 10 hours dark) and temperature (28 °C) in 10 L aquaria, and were fed three times a day *ad libidum* with a commercial diet (GEMMA, Skretting, Wincham, Northwich, UK). Females were bred at weekly intervals to obtain egg batches for CRISPR sgRNA microinjection (MI). The night before spawning, paired males and females bred from different parents were separated by an opaque divider in individual aquaria equipped with marbles at the bottom as the spawning substrate. The divider was removed in the morning, with the fish left undisturbed to spawn. Egg batches in majority containing intact, clean looking, well defined, activated eggs at the 1-cell stage were immediately transferred to microinjection facilities.

For phenotyping observations five couples formed from F3 Hm males and females and five Wt couples were spawned from 3 to 8 times and embryonic development, survival rate, hatching rate, and larval development were subsequently observed until 22 dps. Survival, fecundity and fertilization rate data was collected from 21, 24 and 5 spawns from *vtg1*-KO, *vtg3*-KO and Wt couples, respectively. Hatching rate was calculated based on the number of surviving embryos at 24h and only spawns with > 5 % survival rates were considered, therefore, hatching rate data was collected from 21, 16, and 5 spawns from *vtg1*-KO, *vtg3*-KO and Wt couples, respectively, in this study. Fecundity (number of eggs per spawn) was recorded immediately after spawning and collected eggs were incubated in 100 mm Petri dishes filled with embryo medium (17.1 mM NaCl, 0.4 mM KCl, 0.65 mM MgSO_4_, 0.27 mM CaCl_2_, 0.01 mg/L methylene blue) to assess embryonic development and phenotyping parameters. Incubated eggs/embryos were periodically observed at the early blastula (~256 cell) stage (~2-3 h post spawning hps), at mid-blastula transition stage (~4 hps), at the shield to 75% epiboly stages (~8 hps), at the early pharyngula stage (~24 hps), and during the hatching period at 48 and 72 hps (long-pec to protruding-mouth stages) following standard developmental staging (Kimmel et al., 1995). Fertilization rate was calculated based on viable embryos showing normal cell division and subsequent development to ~24 hps since zygotes failing to initiate cell division, and embryos showing asymmetrical cell cleavage or early developmental arrest were dead by then. As noted above (see Discussion) it is uncertain whether these aberrant eggs/embryos result from infertility or developmental defects. The number of surviving eggs/embryos was recorded, those not surviving were removed and the number of abnormal embryos was recorded at each observation point. Hatched embryos were transferred into larger volume containers (1 L) filled with standard 28°C culture water and were fed *ad libitum* with artemia and GEMMA weaning diet mix after yolk sac absorption (at around 10 dps). At the time of feeding, larvae were also observed for motor and feeding activities. Observations were made daily up to 22 dps. Subsamples of 10-12 embryos and larvae from each clutch were taken for measurements of embryo and chorion diameter, and larval size at 2-3 hps and 8 dps, respectively. Measurements were made using an ocular micrometer under a Zeiss Stemi 2000-C stereomicroscope connected to a ToupCam 3,1 M pixels camera employing the Toupview software.

### 4.2. Single guide RNA (sgRNA) design, synthesis and microinjection

Genomic DNA sequences from all five type-I zebrafish *vtgs* were aligned and three target sites common to all five genes were designed using CRISPR MultiTargeter (Prykhozhij et al., 2015) available online at http://www.multicrispr.net. Of proposed candidates, three target regions located on exons 4, 14 and 17, corresponding to the LvH yolk protein domain were chosen for the *vtg1*-KO experiment. The *vtg3* genomic region was separately submitted to online available target designer tool at http://zifit.partners.org/ZiFiT/ChoiceMenu.aspx (Sander et al., 2007, 2010) and of proposed candidates, three gene specific target regions located on exons 4, 6 and 11, corresponding to the LvH yolk protein domain, were chosen. A schematic representation of the general strategy followed for CRISPR target design is presented in **Fig 1**. Forward and reverse oligonucleotides matching the chosen target sequences (given in **S1 Table**) were annealed and ligated to the pDR274 expression vector (Addgene). The vector was subsequently linearized by the DraI restriction digestion enzyme (Promega) and *in vitro* transcribed using mMessage mMachine T7 Transcription Kit (Ambion) according instructions from the manufacturer. The pCS2-nCas9n plasmid (Addgene Plasmid 47929) was digested with NotI restriction digestion enzyme (Promega) and transcribed using mMessage mMachine SP6 Transcription Kit (Ambion) according instructions from the manufacturer. The sgRNA concentration was measured on a Nanodrop 1000 Spectrophotometer (Thermo Scientific, USA) and integrity was tested before use using an Agilent RNA 6000 Nano Kit (Agilent) on an Agilent 2100 Bioanalyzer.

Approximately 100 eggs per batch were injected with sgRNA mix containing sgRNAs for three target sites (at ~30 ng/ul (=30 mM) in 20ul of the final mix each) and nCas9n RNA (at ~200 ng/ul (=200 mM) in the final mix) at the one-cell stage in both the *vtg1*-KO and *vtg3*-KO experiments. A total of 120 pg sgRNA mix and ~800 pg Cas9 RNA was injected per embryo. Injected embryos were kept in 100 mm petri dishes filled with embryo medium (17.1 mM NaCl, 0.4 mM KCl, 0.65 mM MgSO_4_, 0.27 mM CaCl_2_, 0.01 mg/L methylene blue) to assess microinjection efficiency, embryo survival and development post injection.

### 4.3. Genotyping by conventional PCR

As representatives of their generation, ten embryos were sampled randomly and gDNA was extracted individually and used as a template in targeted conventional PCR reactions to screen for introduced mutations in the targeted *vtg* genes. For this purpose, embryos surviving for 24 h post-injection were incubated in 100 µl of 5 % Chelex^®^ 100 Molecular Biology Grade Resin (BioRad) and 50 µl of Proteinase K Solution (20 mg/ml, Ambion) initially for 2h at 55 °C and subsequently for 10 min 99 °C with constant agitation at 12000 rpm. Extracts were then centrifuged at 5000 xg for 10 minutes and supernatant containing gDNA was transferred into new tubes and stored at −20°C until use.

To evaluate generational transfer of introduced mutations, genotyping of ~2 month old offspring was conducted after extraction of gDNA from fin-clips. For this purpose, fish were anaesthetized in 2-phenoxyethanol (0.5 ml/L) and part of their caudal fin was excised with a sterile scalpel. Genomic DNA from fin tissues were then extracted using Chelex 5 % as described above.

One µl (~ 100 ng) of extracted gDNA was used in 20 µl PCR reactions using AccuPrime™ Taq DNA Polymerase, High Fidelity (Invitrogen) and 10x AccuPrime™ PCR Buffer II in combination with gene specific primers (at 10 µM each) anchoring target sites on the genomic sequence of targeted genes (**Fig 1**). PCR cycling conditions were as follows; 1 cycle of initial denaturation at 94 °C for 2 min, 35 cycles of denaturation at 94 °C for 15–30 sec, annealing at 52–64 °C for 15–30 sec and extension at 68 °C for 1 min per kb plus 1 cycle of final extension at 68 °C for 5 min. Non-purified PCR products or gel purified DNA were sequenced using gene specific primers indicated in **S1 Table** by the Eurofins Genomics sequencing service (https://www.eurofinsgenomics.eu/). Obtained sequences were aligned to corresponding zebrafish genomic sequence using Clustal Omega (Sievers et al., 2011) for characterization and localization of introduced mutations, and then were blasted against all sequences available online using NCBI nucleotide Blast (Blastn) (Altschul et al., 1990) for confirmation of the consistency, accuracy and type of the mutations created at the target sites.

### 4.4. Generation of pure zebrafish lines carrying the introduced mutations

In both the *vtg1*-KO and *vtg3*-KO experiments, embryos carrying introduced mutations were raised to adulthood, fin clipped and re-genotyped to confirm mutation of their type-I or III *vtgs,* and then heterozygous (Ht; *vtg1*+/- and *vtg3+/-)* males with the mutation on a single allele were outcrossed with non-related wild type (Wt; *vtg1*+/+ and *vtg3*+/+) females with no genomic disturbance to produce the F1 generation. Embryos from F1 generation were genotyped as stated above and remaining embryos were raised to adulthood. F1 offspring were screened again at ~2 months of age and, since mutation transmission occurred in two males only per group, these Ht males were crossed with Wt females to produce the F2 generation. Following the same genotyping strategy, F2 Ht males were crossed with Ht females to produce the F3 generation. Finally, F3 homozygous (Hm; *vtg1*-/- and *vtg3*-/-) males and Hm females with both alleles carrying the desired mutation were crossed to produce the F4 *vtg* mutants.

### 4.5. Tissue sampling and analyses

Liver and ovary samples from *vtg1*-KO and *vtg3*-KO F3 Hm, Ht, wt and Wt female zebrafish were excised within 2-3 h after egg collection at the end of phenotyping experiment and after the fish were euthanized with a lethal dose of 2-phenoxyethanol (0.5 ml/L). Ovary samples were aliquoted into four pieces and stored according to subsequent analytical procedures; snap frozen for RNA and protein extraction or placed in Bouin’s solution for histological analyses. Liver samples were aliquoted in two pieces and snap frozen until being used for LC-MS/MS or Western blotting.

### 4.6. Quantitative real time PCR

Total RNA was extracted from frozen liver using TriReagent (SIGMA) and cDNA was synthesized using SuperScript III reverse transcriptase (Invitrogen, USA) from 1 µg of total RNA according to the manufacturer’s instructions. Relative expression levels for all zebrafish *vtgs* (*vtg1, 2, 3, 4, 5, 6* and *7*) in *vtg1*-KO female liver were measured using TaqMan real-time quantitative PCR (RT-qPCR) using gene specific primers and dual-labeled probes (FAM, 6-carboxyfluorescein and a BHQ-1, Black Hole Quencher 1 on 5’ and 3’ terminus, respectively). Sequences of these primers and probes used in this experiment are given in **S1 Table**. Each qPCR was performed in 10 µl reactions containing cDNA (diluted at 1:25), 600 nM of each primer, 400 nM of hybrolysis probe and 1× TaqMan Fast Advanced Master Mix (Applied Biosystems) according the manufacturer’s instructions on a StepOnePlus real time PCR instrument (Applied Biosystems). PCR cycling conditions were as follows: 95°C for 20 seconds, 40 cycles at 95°C for 1 second followed by an annealing-extension at 60°C for 20 seconds. The relative abundance of the target cDNA within a sample set was calculated from a serial dilution curve made from the cDNA pool, using StepOne software (Applied Biosystems). The 2^-ΔΔCT^ mean relative quantification of gene expression method with zebrafish *18S* as a reference gene was employed in this study. Relative expression levels of all zebrafish *vtgs* in *vtg3*-KO female liver were measured using SYBR GREEN qPCR Master Mix (SYBR Green Master Mix kit; Applied Biosystems) as indicated by the manufacturer in a total volume of 10 µl, containing RT products diluted at 1:1000 and 400 nM of each primer in order to obtain PCR efficiency between 95 and 100 %. Sequences of primers used in this experiment are given in **S1 Table**. The RT-qPCR cycling protocol included 3 min initial denaturation at 95 °C followed by 40 cycles of 95 °C for 3 sec and 60 °C for 30 sec on a StepOnePlus thermocycler (Applied Biosystem). The relative abundance of target cDNA within a sample set was calculated from a serially diluted cDNA pool (standard curve) using Applied Biosystem StepOne V.2.0 software. Similarly, the 2^-ΔΔCT^ mean relative quantification of gene expression method with the mean expression value of zebrafish elongation factor 1a (*eif1a*), ribosomal protein 13a (*rpl13a*) and *18S* as reference were employed in this study. Primer sequences and properties for these genes are also given in **S1 Table**. Obtained data was subjected to independent samples Kruskal-Wallis nonparametric test (p<0.05) followed by Benjamini Hochberg correction for multiple tests (p<0.1) (IBM SPSS Statistics Version 19.0.0, Armonk, NY).

### 4.7. Western Blotting

Samples of zebrafish liver, ovary and eggs were homogenized in 100µl of protein binding buffer containing 1mM AEBSF, 10mM Leupeptin, 1mM EDTA and 0.5 mM DTT as indicated by Hiramatsu et al. (2002) using a procellys tissue homogenizer (Bertin Instruments, France). Protein extracts were separated from homogenates with centrifugation at 13 000 rpm +4 °C for 30 minutes to generate supernatant samples for SDS-PAGE. Protein concentrations of the samples were estimated by Bradford Assay (Bradford, 1976) (Bio-Rad, Marnes-la-Coquette, France) and they were diluted to 4 µg protein µl^−1^ in ultrapure water, mixed 1:1 v/v with Laemmli sample buffer (Laemmli, 1970) containing 2-mercaptoethanol, and boiled for 5 min before electrophoresis. A total of 10 µg of sample protein was loaded onto a precast 4–15 % acrylamide gradient Tris–HCl Ready Gel^®^ (BioRad, Hercules, CA) with 4 % acrylamide stacking gel and electrophoresed at 150 V for 45 min using a Tris–glycine buffer system (Laemmli, 1970). Biotinylated protein molecular weight markers (Vector Laboratories, USA) were used to estimate the mass of separated proteins.

Proteins in the gels were transferred to PVDF membranes using a Trans-Blot^®^ Turbo™ Transfer Starter System (BioRad) at 25 mA for 15 min. Blots were blocked for 2 h with Casein solution in tris buffered saline (10 mM Tris HCl containing 15 mM NaCl) and 0.05% Tween 20 (TBST) to reduce non-specific reactions. Affinity purified polyclonal primary antibody raised against a specific peptide epitope on lipovitellin light chain of zebrafish Vtg3 (anti-zfLvL3, GeneScript Custom Antibody production Service, USA) was employed to detect Vtg3 or its product yolk proteins in liver, ovary and eggs from F3 *vtg3*-KO zebrafish. For this purpose, blots were incubated for 2 h at room temperature with the anti-zfLvL3 at a 1:000 dilution in phosphate buffered saline (10 mM Na_2_HPO_4_, pH 7.5, 150 mM NaCl). Membranes were washed three times for 5 minutes in TBST solution and incubated in biotinylated goat anti-rabbit IgG affinity purified secondary antibody diluted 1:8000 in casein solution for 30 minutes at room temperature. Membranes were washed in TBST solution three times for 5 min each and incubated in VECTASTAIN^®^ ABC-AmP™ reagent (VECTASTAIN ABC-AmP Kit, for Rabbit IgG, Chemiluminescent Western Blot Detection, Vector Laboratories) for 10 minutes at room temperature. Following three washes of 5 minutes in TBST, membranes were equilibrated in 0.1 M Tris buffer, pH 9.5 before development in DuoLuX™ Substrate (Vector Laboratories) and exposure to chemiluminescent signal detection on FUSION-FX7 advanced chemiluminescence/fluorescence system (Vilber Lourmat, Germany).

### 4.8. Liquid Chromatography Tandem Mass Spectrometry

Protein extraction of liver and egg samples from *vtg1*-KO, *vtg3*-KO and Wt female zebrafish were done as described by Yilmaz et al. (2017). Briefly, samples were subjected to sonication in 20 mM, pH 7.4, HEPES buffer on ice, soluble protein extracts were recovered following centrifugation (15 000 x g) at +4 °C for 30 min and the remaining pellet was re-sonicated in 30 mM Tris / 8 M Urea / 4 % CHAPS buffer on ice. Ultracentrifugation (105,000 xg) of the pooled protein extracts for 1 h at 4 °C was followed by supernatant recovery and determination of the protein concentration by Bradford Assay (Bradford, 1976) (Bio-Rad, Marnes-la-Coquette, France). Samples of extracts were mixed with sample buffer and DTT and denatured at 70 °C for 10 min before being subjected to SDS-PAGE (60 µg protein/sample lane). When protein samples had completely penetrated the stacking gel (~2 minutes at 200 V-400 mA (~23 W)), electropohoresis was stopped and gels were briefly rinsed in MilliQ ultrapure water (Millipore S.A.S., Alsace, France) and then incubated in fixation solution containing 30 % EtOH / 10 % acetic acid / 60 % MilliQ water for 15 min in order to fix proteins on the gel. Gels were then washed in MilliQ water three times for 5 min each and incubated in EZBlue™ Gel Staining Reagent (Sigma-Aldrich, Saint-Quentin Fallavier, France) at room temperature with slight agitation for 2 h, and de-stained in MilliQ water at room temperature overnight. Subsequently, protein bands were excised from the gel and the excised gel pieces were processed for tryptic digestion and peptide extraction as indicated by Yilmaz et al. (2017). Once peptide extraction was completed, pellets containing digested peptides were resolubilized in 30 µl of 95 % H_2_O : 5 % formic acid by vortex mixing for 10 min and diluted 10 times before being subjected to LC-MS/MS.

Peptide mixtures were analyzed using a nanoflow high-performance liquid chromatography (HPLC) system (LC Packings Ultimate 3000, Thermo Fisher Scientific, Courtaboeuf, France) connected to a hybrid LTQ-OrbiTrap XL spectrometer (Thermo Fisher Scientific) equipped with a nanoelectrospray ion source (New Objective), as previously described (Lavigne et al., 2012; Yilmaz et al., 2017). The spectra search, protein identification, quantification by spectral counts, and spectral count normalization were conducted as described by Yilmaz et al. (2017). To detect significant differences between group mean N-SC values (*vtg1*-KO vs Wt or *vtg3*-KO vs Wt) for different zebrafish Vtgs from liver and eggs, an independent samples Kruskal-Wallis nonparametric test (p<0.05) followed by Benjamini Hochberg correction for multiple tests (p<0.1) was used (IBM SPSS Statistics Version 19.0.0, Armonk, NY).

### 4.9. Ethical Statement

All experiments complied with French & European regulations ensuring ‘animal welfare’ and that ‘Animals will be held in the INRA UR1037 LPGP fish facility (DDCSPP approval # B35-238-6).’ Experimental protocols involving animals were approved by the Comité Rennais d’éthique pour l’expérimentation animale (CREEA).

## 5. COMPETING INTERESTS

Authors declare no competing interests.

## 6. FUNDING

This study was supported by the Region Bretagne in France (SAD-2013)-FishEgg (8210); Project #13009218), the EC-Marie Skłodowska-Curie Actions within the frame of the IEF program (FP7-PEOPLE-2013-IEF; FISHEGG: Project # 626272) and Maternal Legacy (ANR-13-BSV7-0015).

## 7. ACKNOWLEDGMENTS

Authors would like to thank Dr. Amaury Herpin and Dr. Amine Bouchareb for their advices in development of methodological strategies, and Dr. Craig V. Sullivan for their valuable contribution in reviewing and evaluating the manuscript.

## 10. SUPPORTING INFORMATION LEGENDS

**S1 Fig.**
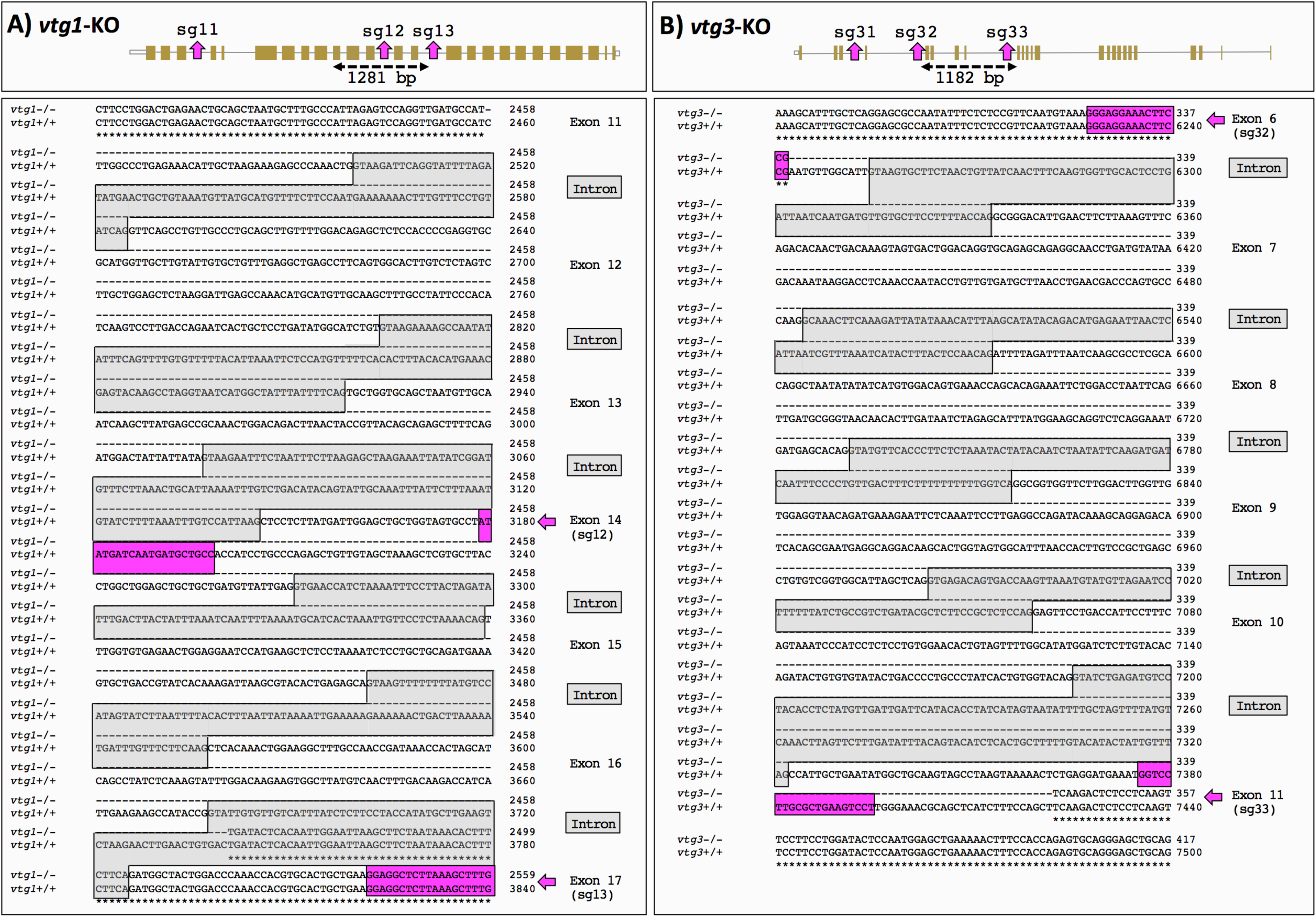
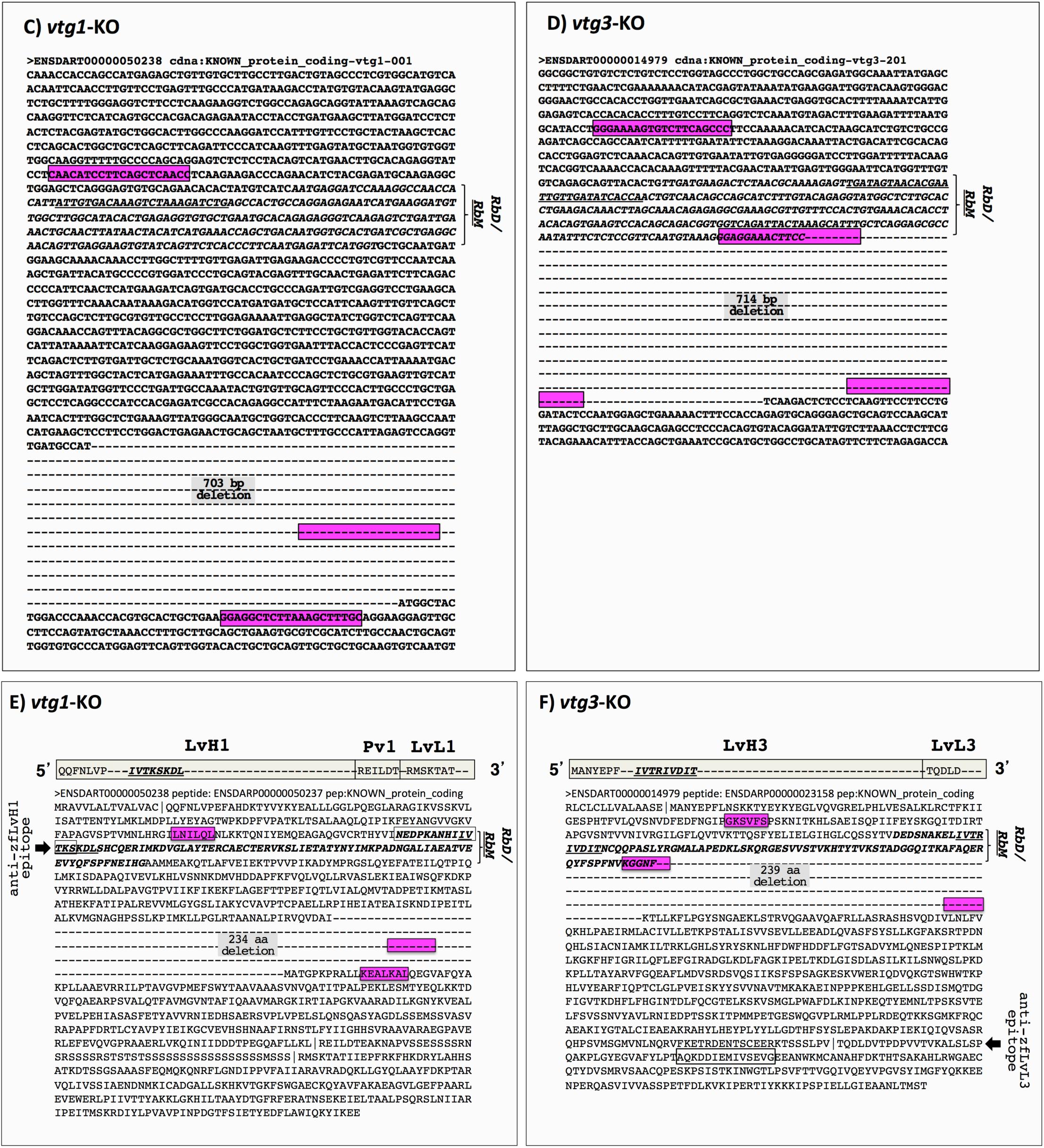
Location and character of mutations introduced by CRISPR/Cas9 in zebrafish *vtgs*. **A-B)** Location on genomic DNA. Schematic representations of the intron/exon structure of zebrafish *vtg1* (representative of type-I *vtgs*) and *vtg3* are given at the top of panels **A** and **B,** respectively. Horizontal line segments indicate introns and filled gold boxes indicate exons. Exons bearing CRISPR/Cas9 target sequences are indicated by magenta-colored arrows pointing upwards to the target name (sg11, sg12, and sg13 for *vtg1;* sg31, sg32, and sg33 for *vtg3).* Horizontal dashed lines bearing dual arrowheads indicate regions where mutations were introduced, with the size of deletions in bp given below the arrows (1281 bp and 1182 bp for *vtg1*-KO and *vtg3*-KO, respectively). The lower sections of panels **A** and **B** show Clustal Omega alignments for partial genomic sequences of the *vtg1* and *vtg3* genes, respectively, covering regions where Cas9 introduced targeted mutations. Sequences of undisturbed wild type alleles are labeled *vtg1*+/+ and *vtg3+/+,* and sequences of homozygous mutated alleles are labeled *vtg1*-/- and *vtg3-/-,* respectively. Dashes were introduced to illustrate regions where deletions occurred in the *vtg1*-/- and *vtg3*-/-sequences. Nucleotide positions are indicated by numbers on the right and asterisks indicate nucleotide identity. Target sequences are enclosed in magenta-colored boxes emphasized by magenta-colored arrows on the right. Intron sequences are given in dark gray font enclosed in light gray filled frames and are labeled by Intron on the right with the same formatting. Exons are shown in regular black font and labeled on the right with exon numbers (e.g. Exon 6, 7, 8…). Exons bearing the target sites are also labeled with the target name below in parenthesis (e.g. Exon 14/(sg12)). **C-D)** Location on predicted cDNA. Nucleotide sequences targeted by sgRNAs for Cas9 editing and present in the predicted transcript are framed in magenta-shaded boxes. The deleted region of the transcript is indicated by dashes replacing nucleotide residues and the size of the deletion in bp is given by gray highlighted text in this region (703 bp deletion for *vtg1* and 714 bp deletion for *vtg3).* The sequence encoding the receptor-binding domain ***(RbD)*** on the LvH of the respective Vtg is shown in italic bold typeface with the sequence encoding the critical, short receptor-binding motif ***(RbM)*** being additionally underlined. **E-F)** Location on predicted polypeptide sequences. Schematic representations of the yolk protein domain structures of Vtg1 (representative of zebrafish type-I Vtgs) and Vtg3 are given in 5’ > 3’ orientation above each panel. Light gray horizontal bars represent the lipovitellin heavy and light chain (LvH, LvL) and the phosvitin (Pv) yolk protein domains of the respective Vtg (Vtg3 lacks a Pv domain) and are labeled above in large bold type. Sequences within these bars indicate the N-terminus of each yolk protein domain, the start of which is also indicated by vertical bars in the polypeptide sequence shown below. The ***RbM*** is shown in bold italic underlined font on the gray horizontal bars in the LvH1 and LvH3 domains. The ***RbD*** and ***RbM*** are also indicated in the polypeptide sequences shown below by bold italic font with the ***RbM*** being additionally underlined. Residues encoded by nucleotide sequences targeted by sgRNAs for Cas9 editing are framed in magenta-shaded boxes. Cas9 created mutations (large deletions) are indicated with dashes replacing amino acid (aa) residues and the size of deletions in aa (234 aa and 239 aa for *vtg1*-KO and *vtg3*-KO, respectively) in these regions are labeled by gray shaded text. Short sequences that were employed as epitopes to develop Vtg domain-specific antibodies against Vtg1-LvH (anti-zfLvH1) and Vtg3-LvL (anti-zfLvL3) are indicated by framed text on the LvH and LvL domains of Vtg1 and Vtg3, respectively, with their location also highlighted by black arrows labeled with the epitope names given by vertically-oriented text in the panel margins.

**S1 Table.**
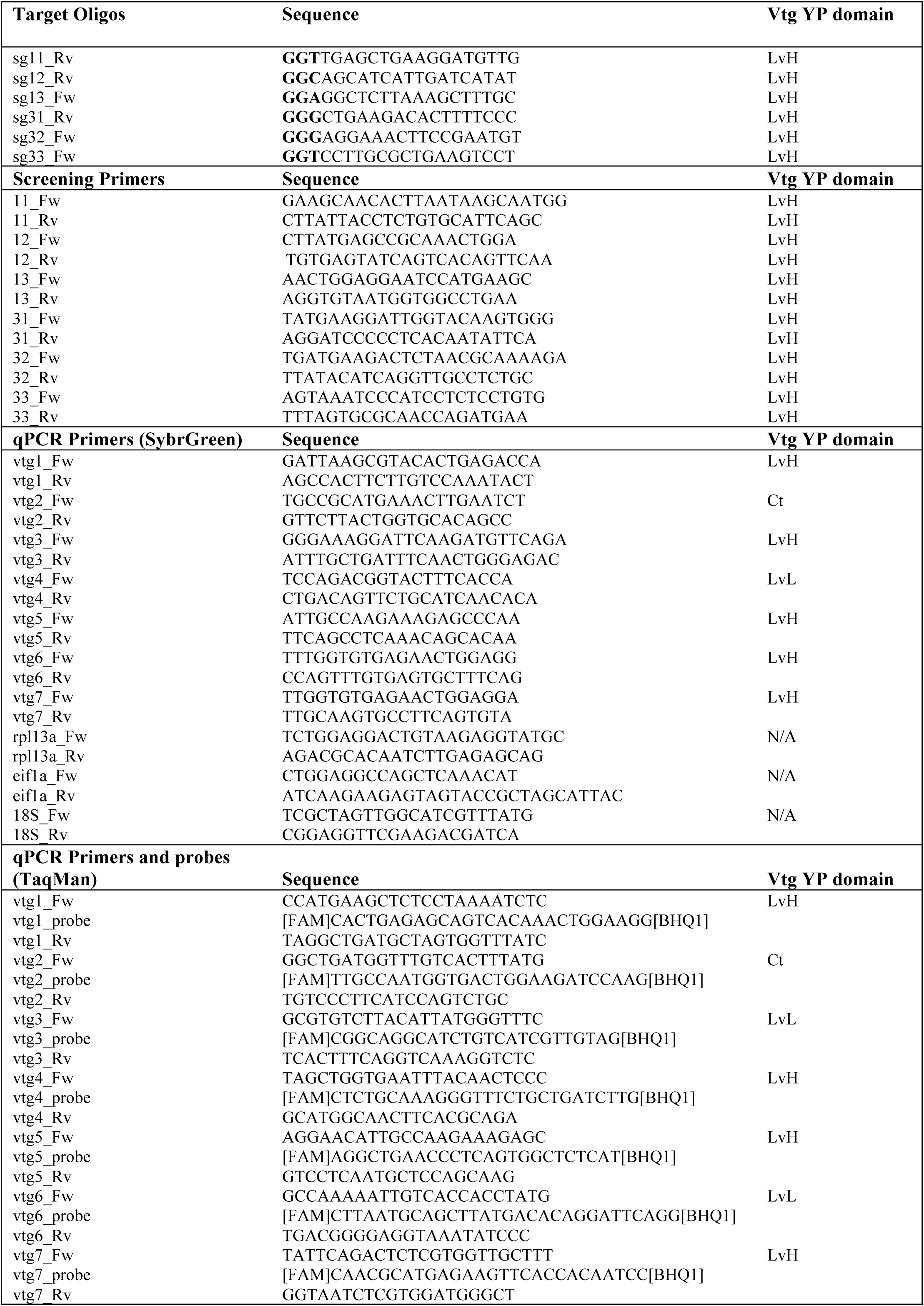
Targets, primers and probes utilized in *vtg1*-KO and *vtg3*-KO studies. Target oligo an screening primer names are given according Figure 1. CRISPR recognition NGG motifs are highlighted by bold typeface on sequences. Position of primers, target sites and probes on vitellogenin (Vtg) yolk protein (YP) domains are given on the far right columns.

